# Disruption of G3BP1 Granules Promotes Mammalian CNS and PNS Axon Regeneration

**DOI:** 10.1101/2024.06.07.597743

**Authors:** Pabitra K. Sahoo, Manasi Agrawal, Nicholas Hanovice, Patricia Ward, Meghal Desai, Terika P. Smith, HaoMin SiMa, Jennifer N. Dulin, Lauren S. Vaughn, Mark Tuszynski, Kristy Welshhans, Larry Benowitz, Arthur English, John D. Houle, Jeffery L. Twiss

## Abstract

Depletion or inhibition of core stress granule proteins, G3BP1 in mammals and TIAR-2 in *C. elegans*, increases axon regeneration in injured neurons, showing spontaneous regeneration. Inhibition of G3BP1 by expression of its acidic or ‘B-domain’ accelerates axon regeneration after nerve injury, bringing a potential therapeutic intervention to promote neural repair in the peripheral nervous system. Here, we asked if G3BP1 inhibition is a viable strategy to promote regeneration in injured mammalian central nervous system where axons do not regenerate spontaneously. G3BP1 B-domain expression was found to promote axon regeneration in the transected spinal cord provided with a permissive peripheral nerve graft (PNG) as well as in crushed optic nerve. Moreover, a cell-permeable peptide (CPP) to a subregion of B-domain (rodent G3BP1 amino acids 190-208) accelerated axon regeneration after peripheral nerve injury and promoted regrowth of reticulospinal axons into the distal transected spinal cord through a bridging PNG. G3BP1 CPP promoted axon growth from rodent and human neurons cultured on permissive substrates, and this function required alternating Glu/Asp-Pro repeats that impart a unique predicted tertiary structure. The G3BP1 CPP disassembles axonal G3BP1, G3BP2, and FMRP, but not FXR1, granules and selectively increases axonal protein synthesis in cortical neurons. These studies identify G3BP1 granules as a key regulator of axon growth in CNS neurons and demonstrate that disassembly of these granules promotes retinal axon regeneration in injured optic nerve and reticulospinal axon elongation into permissive environments after CNS injury. This work highlights G3BP1 granule disassembly as a potential therapeutic strategy for enhancing axon growth and neural repair.

**SIGNIFICANCE STATEMENT:** The central nervous system (CNS) axon does not have the capacity for spontaneous axon regeneration, as seen in the peripheral nervous system (PNS). We previously showed that stress granule-like aggregates of G3BP1 are present in uninjured PNS axons, and these slow nerve regeneration. We now report that CNS axons contain G3BP1 granules, and G3BP1 granule disassembling strategies promote axon regeneration in the injured sciatic nerve, transected spinal cord with a peripheral nerve graft, and injured optic nerve. Thus, G3BP1 granules are a barrier to axon regeneration and can be targeted for stimulating neural repair following traumatic injury, including in the regeneration refractory CNS.

## INTRODUCTION

Though the mammalian peripheral nervous system (PNS) can spontaneously regenerate injured axons, the growth rates are extremely slow at about 1-4 mm/day (1). Regenerating PNS axons can successfully navigate to their targets over short distances and restore at least partial function. However, PNS nerve regeneration for distances more than 5-6 cm is much less successful; this has been attributed to a progressive decline of the growth-supportive environment of the distal nerve and/or the receptivity of target tissues for reinnervation (1). Successful axon regeneration in the central nervous system (CNS) is even more problematic, as CNS neurons have a considerably lower intrinsic growth capacity than PNS neurons and the extracellular environment of the injured CNS actively inhibits axon regeneration (2). Thus, there are unmet clinical needs to accelerate PNS axon regeneration and to enable CNS axon regeneration.

Activation of the mTOR pathway, which increases overall cap-dependent protein synthesis, has been shown to promote axon regeneration in the injured CNS and PNS (3-9). Indeed, deletion or inhibition of PTEN that increases activity of the mTOR pathway promotes regeneration of corticospinal tract axons in the injured spinal cord that are typically among the most refractory to regeneration (7). Protein synthesis in axons has been shown to facilitate PNS axon regeneration (10). Intra-axonal protein synthesis is a well-established mechanism in cultured CNS neurons, where the axon uses locally synthesized proteins for directional growth; there are now multiple lines of evidence indicating that many CNS neurons retain the capacity for intra-axonal protein synthesis well into adulthood (11-13), and some evidence for mRNA translation in spinal cord axons (14-17). We previously showed that axonal RNA-protein granules containing the core stress granule protein G3BP1 slow PNS axon regeneration by sequestering mRNAs and attenuating their translation in injured axons (18). Stress granules, which form by liquid-liquid phase separation (LLPS), are typically seen during metabolic or oxidative stress conditions and are used to store mRNAs encoding proteins that are not needed to respond to the stress (19). However, axons use stress granule-like structures as RNA storage depots in the apparent absence of any stress signals (18, 20), and disassembly of axonal G3BP1 granules promotes intra-axonal protein synthesis and accelerates PNS axon regeneration (18, 20). Here, we asked whether inhibiting the RNA sequestration function of G3BP1 has therapeutic potential for promoting CNS axon regeneration.

## RESULTS

### G3BP1 granules attenuate axon regeneration in the central nervous system

We previously showed that G3BP1 granules slow PNS axon regeneration and that expressing G3BP1’s acidic domain or ‘B-domain’, consisting of rat G3BP1 amino acids 141-220 (UniProt ID: D3ZYS7_RAT), accelerates axon regeneration following sciatic nerve crush injury (18). G3BP1 B-domain expression decreases both the size and number of axonal G3BP1 granules, increases intra-axonal protein synthesis in cultured sensory neurons, and accelerates PNS nerve regeneration (18). Thus, the exogenously expressed G3BP1 B-domain acts as a ‘dominant negative’ agent. With these effects seen in PNS axons when G3BP1 is inhibited, we hypothesized that axonal G3BP1 granules could also impede CNS axon regeneration.

Neural stem cells have been shown to promote CNS axon growth, where injured corticospinal axons can regrow to form a relay circuit across a site of spinal cord injury by forming synapses on stem cell-derived neurons grafted into the lesion site (21, 22). Lu *et al*. (2012) showed that neural progenitor cells isolated from the embryonic spinal cord (‘caudalized grafts’) promote growth of host corticospinal motor axons, whereas neural progenitor cells isolated from the embryonic telencephalon (‘rostralized grafts’) do not support regeneration into the graft (21). Using immunofluorescence for G3BP1 and TIA1, another core stress granule protein, we find that corticospinal axons in injured spinal cord contain more G3BP1-TIA1 granules in animals with a rostralized progenitor graft, which does not support regeneration, than those with a caudalized progenitor graft (**Fig. 1A**). This emphasizes that spinal cord axons contain G3BP1 and TIA1 granules, and that the abundance of these intra-axonal stress granule-like structures relatively decreases when a regeneration-promoting stimulus is provided to the injured spinal cord axons.

**Figure 1:**
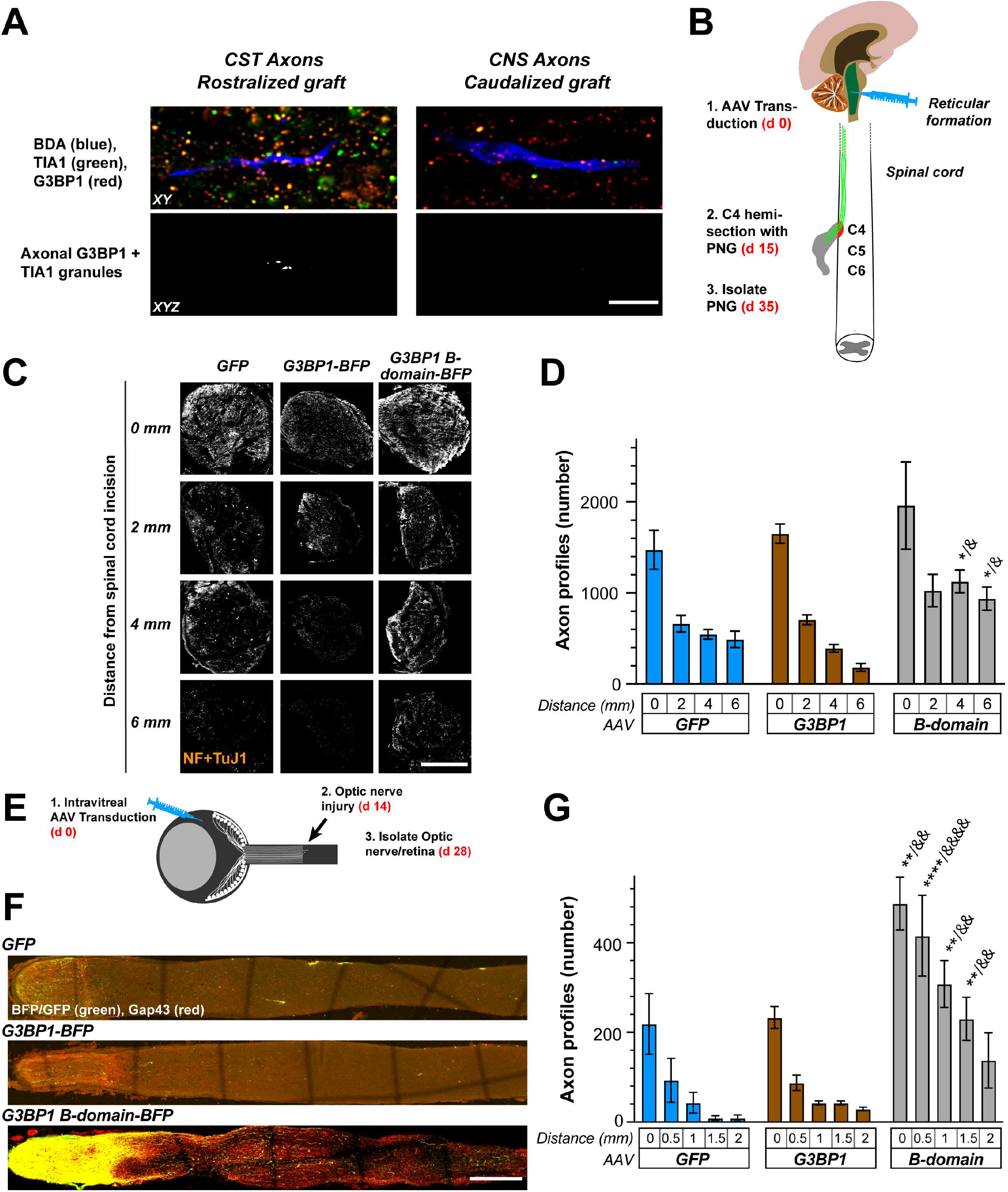
Expression of G3BP1 acidic domain in CNS neurons facilitates axon regeneration. ***A***, Representative exposure-matched confocal images of corticospinal axons in spinal cord labeled with biotinylated dextran amine (BDA) following C4 level contusion injury and subsequent grafting with rostralized or caudalized stem cell graft (21). Upper row shows XY optical plane through CST axon regenerating into graft; lower row shows signals for stress granule proteins G3BP1 and TIA1 that overlap with BDA across Z sections as XYZ projection in a separate channel (‘axonal G3BP1 + TIA1‘). There are stress granules present in the few axons regenerating into rostralized grafts, and these are absent in the more numerous axons regenerating into caudalized grafts [scale bar = 10 µm]. ***B***, Schematic of experimental paradigm and timeline for spinal cord injury model used in C-D, with regeneration of reticulospinal neurons in the permissive environment of a peripheral nerve graft (PNG). ***C***, Exposure-matched epifluorescence images for NF-+ βIII Tubulin-immunostained PNGs from animals transduced with AAV5-GFP, -G3BP1-BFP, or -G3BP1 B domain-BFP shown at indicated distances from grafting site at proximal side of transected spinal cord. For G3BP1 granule size and density after CPP treatment see Supplemental Figure S1A-C [scale bar = 500 µm]. ***D***, Quantitation of regenerating reticulospinal axon profile numbers in PNGs from panel C at indicated distances away from grafting site in spinal cord shown as mean ± standard error of the mean (SEM; N = 5 animals; *p⍰≤⍰0.05 vs. GFP- and & p⍰≤⍰0.05 vs. G3BP1-BFP by two-way ANOVA with Tukey HSD post-hoc). ***E***, Schematic of experimental paradigm for AAV transduction of retinal ganglion cells (RGC) and optic nerve crush injury used in F-G. ***F***, Confocal montage images of optic nerves for AAV2-GFP, -G3BP1-BFP, G3BP1 B-domain-BFP transduced animals at 14 days post injury. Regenerating axons are detected by anti-GAP43 (red) and GFP, G3BP1-BFP and G3BP1 B-domain-BFP signals by anti-BFP/GFP antibodies (green). For G3BP1 granule size and density after CPP treatment see Supplemental Figure S1D-F [scale bar = 250 µm]. ***G***, Quantitation of GAP43 signals from F for regenerating RGC axon profiles is shown as mean ± SEM (N ≥ 5 animals; **p⍰≤⍰0.01 and ****p⍰≤⍰0.0001 vs. GFP- and && p ≤ 0.01 and &&&& p ≤ 0.001 vs. G3BP1-BFP-transduced animals by two-way ANOVA with Tukey HSD post-hoc). For RGC survival data see Supplemental Figure S2.

Adult dorsal root ganglion (DRG) sensory neurons and embryonic cortical neurons from rats show increased axon growth upon disassembly of G3BP1 granules induced by expression of the G3BP1 B-domain in culture (18). To test the possibility that G3BP1 granules may negatively influence the growth of adult CNS axons, we asked whether B domain expression would increase rates of injured spinal cord axon growth within a permissive growth environment. For this, we transduced reticulospinal neurons using adeno-associated virus (AAV), serotype 5 expressing G3BP1 B-domain-blue fluorescent protein (BFP), full length G3BP1-BFP, or GFP. Fifteen days later, a C4 hemisection was performed and a peripheral nerve graft (PNG) was apposed to the rostral injury site adjacent to the reticulospinal tract as a growth-permissive environment (**Fig. 1B**). Immunofluorescence for axons performed at 20 days after spinal cord injury/PNG placement showed more axons extending further distances from the spinal cord/PNG apposition in the G3BP1 B-domain-BFP expressing animals compared to the G3BP1-BFP and GFP expressing animals (**Fig. 1C-D**). These data indicate that G3BP1 B-domain expression can promote adult CNS axon growth *in vivo*.

The axons regenerating in the PNG above provide some similarities for spinal motor axons regenerating in peripheral nerve – *i*.*e*., the descending reticulospinal neurons regenerated their axons through a growth-permissive PNS environment. Thus, we moved to an optic nerve injury model where CNS neurons must regenerate their axons through a growth-inhibitory CNS environment. For this, retinal ganglion cells (RGC) were transduced via intravitreal injection with AAV serotype 2 (AAV2) expressing G3BP1 B-domain-BFP, G3BP1-BFP, or GFP, and then crushed the optic nerve 14 days later (**Fig. 1E**). 14 days after this injury, immunofluorescent staining for growth-associated protein 43 (GAP43) to detect regenerating axons and BFP/GFP to detect transduced neurons showed that G3BP1 B-domain expression in RGCs greatly increases axon regeneration (**Fig. 1F-G**). Interestingly, animals expressing the G3BP1 B-domain appeared to show far higher number of axons proximal to the injury site (**Fig. 1F**). Evaluation of transduction efficiency based on optic nerve BFP positivity showed equivalent axon percentages for G3BP1-BFP vs. G3BP1 B-domain-BFP (**Suppl. Fig. S1A-B**).

Expression of G3BP1 B-domain in PNS neurons disassembles axonal G3BP1 granules and in turn accelerates PNS nerve regeneration (18). To determine if expression of the G3BP1 B-domain affects G3BP1 granules in CNS axons, we compared G3BP1 granules in the reticulospinal axons in PNGs and RGC axons in the injured optic nerve using confocal microscopy where we could extract G3BP1 granule signals that overlapped with axonal markers across optical planes as a separate panel (projected XYZ images shown for ‘Axonal G3BP1’ panels in **Suppl. Fig. S1C and F**). G3BP1 B-domain expression in reticulospinal neurons and RGCs resulted in smaller and fewer G3BP1 granules in the axons compared to G3BP1 and control GFP expression (**Suppl. Fig. S1D-E and G-H**). In contrast to many other CNS and PNS axotomies, optic nerve crush triggers death in the majority of RGCs (23-26), and we find that transduction with AAV2-G3BP1 B-domain-BFP increased post-injury RGC survival compared to the two control conditions (**Suppl. Fig. S2**). Taken together, these findings indicate that expression of the G3BP1 B-domain can facilitate regeneration of injured CNS axons and overcome the growth-inhibitory environment of the injured CNS, likely through G3BP1 granule disassembly and release of axonal mRNAs for translation.

### Cell permeable G3BP1 B-domain peptide increases PNS nerve regeneration

Using synthetic cell-permeable peptides (CPP), we previously found that amino acids 147-166 and 190-208 of rat G3BP1 increase axon growth from cultured sensory and cortical neurons (18), whereas a CPP consisting of rat G3BP1 amino acids 168-189 had no significant effect (18). G3BP1 amino acid sequences for 147-166 and 190-208 CPPs are quite acidic with Glu or Asp residues occupying 11 of 21 (52.4%) and 8 of 19 residues (42.1%), respectively (**Fig. 2A**). The G3BP1 190-208 CPP was more effective than 147-166 CPP in our earlier study (18), so we tested whether the G3BP1 190-208 CPP might affect PNS axon regeneration *in vivo* using a clinically relevant paradigm where the CPP is applied after nerve injury. For this, a mid-thigh sciatic nerve crush was performed on adult rats and 2 days later the G3BP1 190-208 or 168-189 CPP was injected just proximal to the injury site to achieve an estimated 113 µM concentration within the perineurium (**Fig. 2B**). G3BP1 190-208 CPP-injected animals showed a strong increase in axon regeneration at 7 days post-injury (5 days after peptide treatment) compared to G3BP1 168-189 CPP-injected animals, an effect that was particularly evident at longer distances down the nerve (**Fig. 2C-D**). The G3BP1 168-189 CPP-injected animals showed no significant difference from those injected with an equivalent volume of the vehicle control (phosphate-buffered saline [PBS]; **Fig. 2D**).

**Figure 2:**
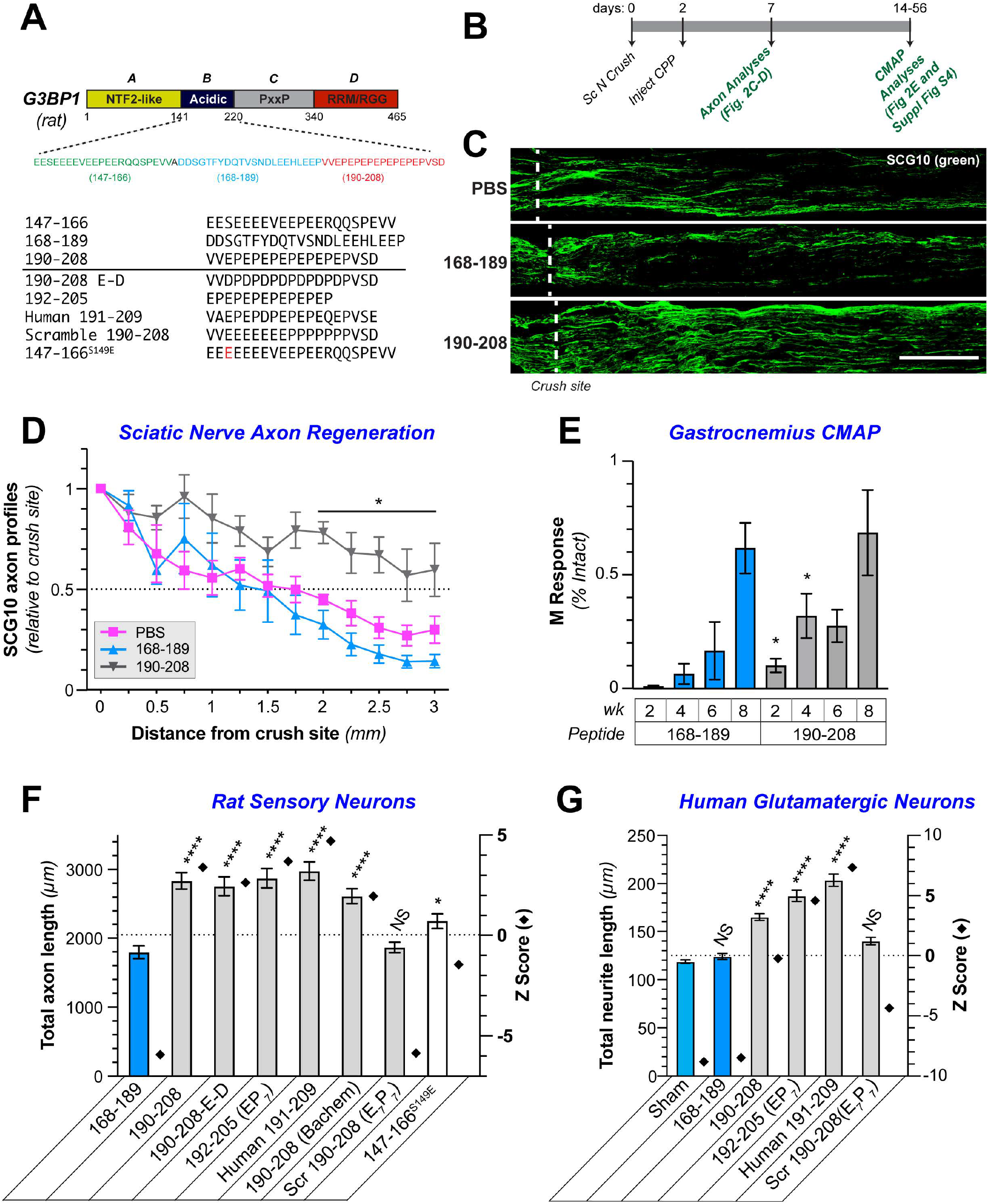
G3BP1 190-208 peptide promotes PNS and CNS axon growth. ***A***, Schematic of rat G3BP1 protein with defined domains shown on the top. Sequence of acidic domain (B domain) of G3BP1 shown below with sequence of amino acids 147-166, 168-189 and 190-208 used for cell permeable peptides (CPP). Variants of these CPPs tested in panels F-G and Supplemental Figures S3-4 are indicated. ***B***, Timeline for sciatic nerve injury followed by intra-nerve G3BP1 CPP treatments used for panels C-E and Supplemental Figure S3A. ***C-D***, Exposure-matched epifluorescence images for anti-SCG10 immunostained sciatic nerves 7 days post-crush that were treated with indicated CPPs or PBS are shown in C. Dotted line shows the injury site. Quantitation of SCG10+ axon profiles at indicated distances past the injury site from C is shown in D as mean ± SEM (N ≥ 6 animals; * p ≤ 0.05 by two-way ANOVA vs. 168-189 with Tukey HSD post-hoc) [scale bar = 500 µm]. ***E***, Compound muscle action potentials (CMAP) are shown for gastrocnemius muscle of animals treated with G3BP1 CPPs after sciatic nerve crush injury as in B are shown as average % intact M responses ± SEM for indicated times (N ≥ 4 animals; *p1≤10.05 by Student’s t-test for indicated data pairs). For CMAP data on tibialis anterior muscle see Supplemental Figure S3A. ***F***, Quantitation of total axon growth of cultured adult rat sensory (DRG) neurons treated with G3BP1 CPP and variants is shown as mean ± SEM (left Y-axis) and Z score relative to population mean (right Y-axis; N ≥ 352 neurons across three biological replicates for each condition; * p ≤ 0.05 and **** p ≤ 0.0001 vs. 168-189 CPP by one-way ANOVA with Dunnett post-hoc). Representative images and data on longest axon per neuron and branch density are shown in Supplemental Figure S3. ***G***, Quantitation of total neurite length for human iPSC (hiPSC)-derived glutamatergic neurons untreated (Sham) or treated with indicated G3BP1 CPPs and variants shown as mean ± SEM (left Y-axis) and Z score vs. population mean (right Y-axis; N ≥ 1165 neurons across three biological replicates for each condition; **** p ≤ 0.0001 vs. Sham by one-way ANOVA with Tukey HSD post-hoc). Representative images plus longest neurite length and neurite branching data are shown in Supplemental Figure S4.

To determine if the accelerated nerve regeneration seen with the G3BP1 190-208 CPP brings functional improvements, we compared neuromuscular junction reinnervation in G3BP1 168-189 vs. 190-208 CPP-treated animals using compound muscle action potentials (CMAP) for the lateral gastrocnemius (LG) and tibialis anterior (TA) muscles (**Fig. 2B**). Recovery of CMAPs was significantly accelerated in the animals treated with G3BP1 190-208 CPP treatment for both the LG and TA at 2 and 4⍰weeks after sciatic nerve crush (**Fig. 2E, Suppl. Fig. S3A**). There were no differences between the G3BP1 190-208 and 168-189 CPP-treated animals at 8 weeks post-injury (**Fig. 2E**), consistent with spontaneous regeneration in the PNS; similar results were previously reported for sciatic nerve crush recovery in AAV-G3BP1 B-domain transduced animals (18). It should be noted that significant differences between the G3BP1 190-208 and 168-189 CPP-treated animals were not seen at 6 weeks post-injury, which points to a limited *in vivo* duration for the G3BP1 190-208 CPP activity and emphasizes an opportunity to improve peptide stability.

The G3BP1 190-208 sequence contains seven Glu-Pro (EP_7_) repeats flanked by N-terminal Val-Val and C-terminal Val-Ser-Asp (**Fig. 2A**). We sought to determine critical residues for function of the 190-208 G3BP1 CPP by asking whether sequence alterations modified its axon growth-promoting activity in adult DRGs cultured on laminin. We initially tested 190-208 G3BP1 CPP from a different vendor (‘Bachem’) and got identical results to the previous synthetic peptide (**Fig. 2F, Suppl. Fig 3B-D**). A modified 190-208 CPP in which all Glu residues in the EP7 region were replaced with similarly acidic Asp residues promoted axon growth to a similar extent as the original 190-208 CPP (**Fig. 2F, Suppl. Fig. S3B-D**). Similarly, a CPP lacking the N- and C-terminal residues flanking the EP7 repeats (G3BP1 192-205 CPP) maintained full axon growth-promoting activity (**Fig. 2F, Suppl. Fig. S3B-D**), indicating that just the seven EP repeats are sufficient to promote axon growth. However, a scrambled 190-208 G3BP1 CPP where the EP7 was changed to seven Glu followed by seven Pro (E7P7) generated axon lengths that were not significantly different than the non-functional 168-189 G3BP1 CPP (**Fig. 2F, Suppl. Fig. S3B-D**). Taken together, these data indicate that axon growth-promoting functions of the 190-208 G3BP1 CPP require repeats of acidic residues with intervening Pro.

The G3BP1 147-166 sequence bears some similarities to the 190-208 sequence with a preponderance of Glu residues (**Fig. 2A**). Interestingly, we have shown that Casein kinase 2 alpha (CK2α)-dependent phosphorylation of G3BP1 on Ser 149 (G3BP1^PS149^) causes G3BP1 granule disassembly and decreases G3BP1’s RNA binding activity (20). Considering the functional effects of Ser 149 phosphorylation, we asked whether replacing Ser 149 with an acidic residue to mimic phosphorylated G3BP1 (147-177^S149E^) would increase the axon growth effects of G3BP1 147-166 CPP to levels comparable to the G3BP1 190-208 CPP. G3BP1 147-149^S149E^ CPP treatment generated axon lengths intermediate between the 168-189 and 190-208 CPPs (**Fig. 2F, Suppl. Fig. S3B-D**), indicating that the S149E modification did not raise the 147-166 CPP’s efficacy to that of the 190-208 CPP. We further tested a CPP containing the human G3BP1 sequence corresponding to rat amino acids 190-208 (human 191-209). The human 191-209 CPP promoted axon growth comparable to the rat 190-208 CPP despite an Asp at residue 197 (instead of Glu) and Gln at residue 204 (instead of Pro) within the EP7 region (**Fig. 2A and F, Suppl. Fig. S3B-D**). Rat 190-208, rat 192-205 (EP_7_), and human 191-209 CPPs also substantially increased neurite growth from glutamatergic neurons derived from human induced pluripotent stem cells (hiPSC) (**Fig. 2G, Suppl. Fig. S4**).

Axon branching in adult DRG cultures treated with the G3BP1 CPP variants was also assessed. The rat 192-205, human 191-209, and rat 147-166^S149E^ G3BP1 CPPs increased branch density by ∼12% as compared to the rat 190-208 and other CPP variants, which were not different than the rat 168-189 CPP with no axon outgrowth effects (**Suppl. Fig. S3D**). Similarly, rat 192-205 and human 191-209 G3BP1 CPPs increased neurite branch density in the hiPSC-derived neurons, but the rat 190-208 and scrambled 190-208 G3BP1 CPPs were no different than the rat 168-189 CPP or no treatment control (**Suppl. Fig. S4C**). Notably, each of the CPPs that increased the total axon/neurite length also increased the length of the longest axon/neurite per neuron (**Suppl. Figs. S3C and S4B**), indicating that the increased branching seen with rat 192-205 and human 191-209 CPPs does not account for the difference in total axon/neurite lengths shown above. In contrast, the increase in DRG total axon length and longest axon per neuron induced by the rat 147-166^S149E^ CPP was modest with non-significant Z score compared to the population mean, suggesting that it had minimal effects on elongating axon growth (**Fig. 2F, Suppl. Fig. S3C-D**).

### G3BP1 190-208 cell-permeable peptide differentially alters CNS axon regeneration depending on the local environment

Since the G3BP1 190-208 CPP successfully promotes PNS axon regeneration (**Fig. 2B-E**) and the G3BP1 B-domain promotes regeneration of reticulospinal axons (**Fig. 1B-D**), we asked whether this CPP strategy might also be effective in promoting CNS axon regeneration. To test this, we performed a C4 hemisection with a PNG apposed close to the rostral injury site for axon growth by descending axons or local interneurons (**Fig. 3A**). The same day, the rat 168-189 or 190-208 G3BP1 CPPs were injected directly into the spinal cord just rostral to the injury site and regeneration through the PNG was evaluated 10 days later by immunostaining for axonal markers (**Fig. 3A**). Increases in axon profile numbers were only seen in the most proximal segment of the PNG for the G3BP1 190-208 compared to 168-189 CPP-treated animals (**Fig. 3B**). Thus, in contrast to the effect of the G3BP1 B-domain in this experimental model, there were no differences between treatment groups in the more distal PNG segments (2-6 mm; **Fig. 3B**). Confocal microscopy for XYZ imaging of the proximal PNG segment was used to reconstruct the path and branches of individual regenerating axons across optical planes representing cross sections of the PNG in combination with *Neurolucida* software neurite tracking/branching function. Surprisingly, the 190-208 G3BP1 CPP-treated animals showed a remarkable increase in axon branching complexity in the proximal PNG compared to the 168-189 CPP-treated animals (**Fig. 3C-D**). Thus, although the 190-208 G3BP1 CPP increases axon outgrowth in PNGs, this appears to result in axon branching or sprouting rather than the axon elongation seen in PNGs with AAV-G3BP1 B-domain transduced reticulospinal neurons.

In the above spinal cord injury experiment, the CPPs were presented to axons directly in the injured spinal cord but the nerve regeneration experiments in Figure 2 presented the CPPs to axons in the permissive environment of the peripheral nerve. Thus, we asked if exposing regenerating reticulospinal axons to G3BP1 CPPs within the PNG might alter their growth differently. For this, we performed a C4 hemisection with descending PNG as in Figure 1; 30 days later, CPPs were injected into the distal PNG that was then apposed to the spinal cord adjacent to the caudal injury site (**Fig. 3E**). Animals were euthanized 22 days later (52 days post-injury, 22 days CPP exposure) and spinal cords with attached PNG were analyzed for axon regeneration using SCG10 immunostaining. The 190-208 G3PB1 CPP-treated animals showed increased numbers of axons entering into the spinal cord from the PNG, and those axons extended considerably greater distances into the spinal cord compared to scrambled 190-208 CPP-treated animals (**Fig. 3F-G**). Thus, the 190-208 G3BP1 CPP can promote growth of injured adult spinal cord axons, but our data suggest that it generates different growth morphologies depending on whether the CPP is presented in the growth-inhibitory environment of the spinal cord or the growth-permissive environment of the PNG.

**Figure 3:**
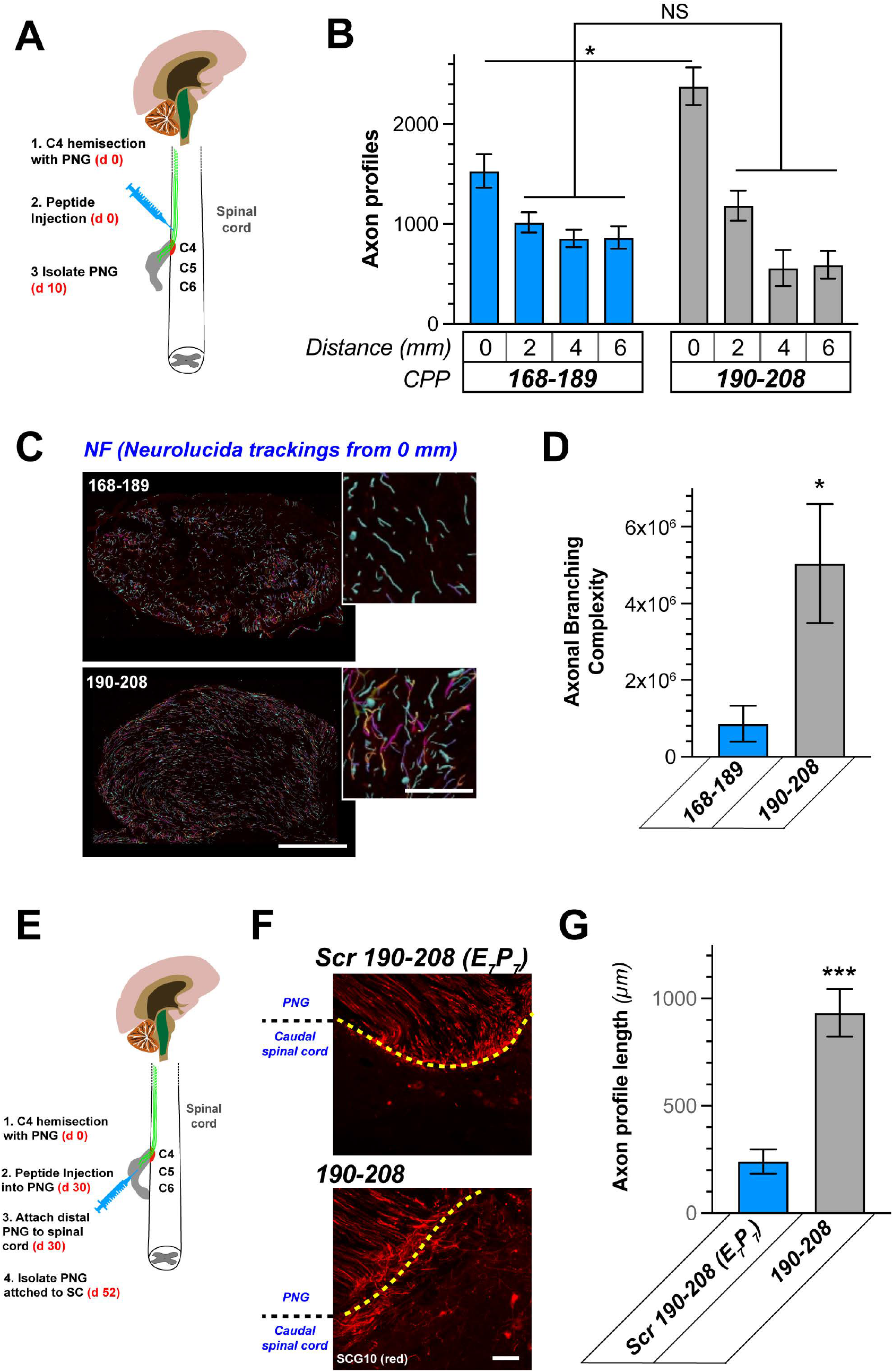
G3BP1 190-208 cell permeable peptide promotes in vivo CNS axon regeneration. ***A***, Schematic of experimental paradigm for spinal cord injury plus PNG and CPP treatment into spinal cord just rostral to injury site used in B-D. ***B***, Quantitation of regenerating CNS axon profiles into PNGs from panel B at indicated distances from spinal cord shown as mean ± SEM (N = 5 animals; *p⍰≤⍰0.05 by Student’s t-test for indicated data pairs). ***C***, Neurolucida tracing images for NF-+ -βIII tubulin-immunostained regenerating axons in proximal PNG shown. Primary branches are pseudo-colored as cyan and secondary/tertiary branches are pseudo-colored from violet-to-magenta. Note increased violet to magenta neurites in the G3BP1 190-208 CPP-vs. 168-189 CPP-treated animals indicating increased branching [scale bar = 250 µm in large images and 25 µm in insets]. ***D***, Quantitation of axonal branching from Neurolucida tracing from C shown as mean ± SEM (N = 5 animals; *p⍰≤⍰0.05 by Student’s t-test). ***E***, Schematic of experimental paradigm for spinal cord injury plus PNG and CPP treatment into the PNG used in F-G is shown. ***F***, Representative fluorescent images show anti-SCG10 immunosignals for regenerating CNS axons in PNGs. Yellow dotted line shows the junction between PNG and spinal cord, with many more axons penetrating the spinal cord in G3BP1 190-208 CPP-than scrambled 190-208 (Scr 190-208 [E7P7])-treated animals [scale bar = 50 µm]. ***G***, Quantitation of distal spinal cord reticulospinal axons from F shown as mean ± SEM (N = 6 animals; ***p⍰≤⍰0.001 by Student’s t-test).

### Axonal growth morphology induced by G3BP1 190-208 cell permeable peptide depends on growth environments

To directly test the possibility that the G3BP1 190-208 CPP differentially alters axon growth depending on the permissiveness of the growth substrate, we evaluated neurite growth from embryonic day 18 (E18) rat cortical neurons cultured on growth-permissive (poly-D-lysine [PDL]) or growth-inhibitory (PDL + aggrecan) substrates. For neurons cultured on PDL, rat 190-208 G3BP1 CPP nearly doubled total axon length and distance the axon extended from the soma, had a smaller but significant effect on dendrite lengths, and decreased axon and dendrite branching compared to 168-189 CPP (**Fig. 4A, Suppl. Fig. S5**). In contrast, 190-208 G3BP1 CPP-treated cortical neurons cultured on PDL + aggrecan showed increased axon and dendrite branching compared to 168-198 CPP-treated neurons (**Fig. 4B**). Total axon and dendrite lengths were increased by 25% and 80%, respectively, for the 190-208 G3BP1 CPP-treated compared to 168-198 CPP-treated cortical neurons cultured on PDL + aggrecan; distance extended from the soma was not changed for the axons but was increased for the dendrites in these CNS neurons (**Fig. 4B, Suppl. Fig. S5B**). We had previously shown that G3BP1 190-208 CPP treatment increases axon lengths for DRG neurons grown on laminin (18). In contrast to the cortical neurons used above, DRG neurons only extend Tau ^+^/MAP2^-^ neurites with other axonal features (27-29) in culture, so only axon growth can be evaluated in these DRG cultures. 190-208 G3BP1 CPP-treated DRG cultures showed increased total axon length on laminin + aggrecan, but axon branching was also significantly increased compared to the 168-189 CPP-treated neurons (**Suppl. Fig. S5C-D**). Taken together, these data indicate that disassembling axonal G3BP1 granules using the 190-208 G3BP1 CPP in non-permissive growth environments increases neurite branching, while the CPP promotes elongating axon growth in permissive growth environments.

**Figure 4:**
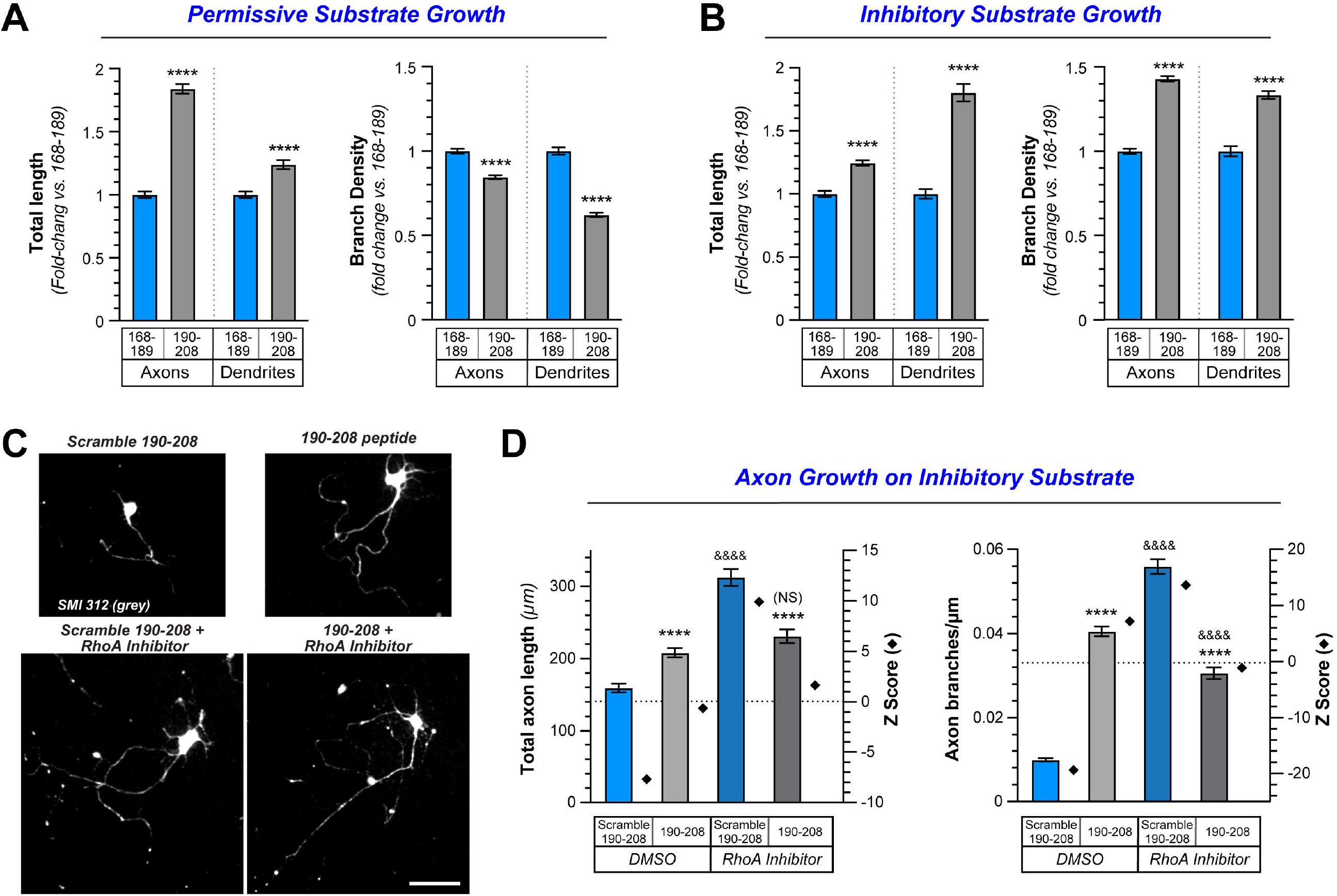
G3BP1 acidic domain inhibits axonal G3BP1 granule assembly to enable CNS axon regeneration. ***A-B***, Axon and dendrite length and branching for G3BP1 190-208 vs. 168-189 CPP-treated E18 rat cortical neurons cultured on growth-permissive (A; PDL) vs. growth-inhibitory (B; PDL + aggrecan) substrates is shown as mean ± SEM. Note that axon and dendrite branching are significantly increased on the growth inhibitory substrates with 190-208 CPP exposure (N ≥ 1042 neurons across 3 culture preparations for each condition; **** p ≤ 0.0001 by students T-test). Axon morphologies for adult DRG cultured on aggrecan plus G3BP1 190-208 vs. 168-189 CPP shown in Supplemental Figure S5C-D. ***C***, Representative epifluorescence images of SMI312 immunostained E18 rat cortical neurons cultured on aggrecan and treated with G3BP1 190-208 CPP vs. scrambled 190-208 CPP ± 10 µM RhoA kinase inhibitor shown [scale bar⍰=⍰50⍰µm]. ***D***, Quantitation of axon length and branching density for E18 cortical neurons as in C shown as mean ± SEM (N ≥ 259 neurons across three repetitions for each condition; ****/&&&& p⍰≤⍰0.0001 by one-way ANOVA with Tukey HSD post-hoc; * for indicated treatment group vs. scrambled 190-208-DMSO, & for indicated treatment group vs. 190-208-DMSO). Average length axons extend beyond soma shown in Supplemental Figure S5E.

Axon growth-inhibiting vs. -promoting substrates are known to differentially regulate activities of the small GTPases RhoA, Rac, and CDC42 (30-34). CSPGs, in particular, activate RhoA in distal axons, which promotes actin filament disassembly and growth cone retraction (2), and accordingly, inhibition of RhoA allows axons to grow on this non-permissive substrate (35-37). In contrast, axon growth-promoting stimuli (*e*.*g*., neurotrophins) activate Rac and CDC42 in distal axons, which promotes actin filament assembly and growth cone advance (34). To determine if RhoA plays any role in the increased axon branching seen with rat 190-208 G3BP1 CPP treatment for neurons grown on the CSPG aggrecan, we pharmacologically inhibited RhoA kinase (ROCK) using the ROCK inhibitor Y27632 (10 µM) in E18 rat cortical neurons cultured on PDL + aggrecan simultaneously with CPP treatment. As Y27632 has been previously shown to increase neurite growth of both PNS and CNS neurons on growth non-permissive substrates (38-40), we focused on a combinatorial treatment of the neurons with Y27632 and CPP. For the control CPP-treated neurons (scrambled 190-208 G3BP1), RhoA inhibition increased total axon length/neuron, but this can be attributed to an approximately five-fold increase in axon branching compared to vehicle-treated neurons (DMSO; **Fig. 4C-D, Suppl. Fig. S5E**). For the rat 190-208 G3BP1 CPP-treated neurons, RhoA inhibition increased total axon length, but decreased axon branching compared to DMSO-treated cultures (**Fig. 4D, Suppl. Fig. S5E**). Together, these data indicate that RhoA activity drives axon branching on the growth-inhibitory substrate in the presence of the G3BP1 granule disassembling CPP. This raises the interesting possibility that intra-axonal signaling, driven by the permissiveness vs. non-permissiveness of the axon’s environment, can shift the elongating morphology of CNS axon growth depending upon RhoA activity and G3BP1 granule formation status.

### G3BP1 192-205 cell permeable peptide disassembles specific axonal RNPs and enhances axonal protein synthesis

Previously, we have shown that the expression of the G3BP1 B-domain in NIH-3T3 cells blocks arsenite-induced SG formation (18). Moreover, the application of the G3BP1 190-208 CPP to DRG neurons leads to immediate disassembly of axonal G3BP1 granules and specifically increases axonal protein synthesis (18). In Figure 2, we identified that the core 192-205 (EP_7_) peptide of rat G3BP1 is equally effective as the 190-208 peptide. Based on these data, we asked if the G3BP1 granule disassembling 192-205 (EP_7_) CPP works in a similar manner in CNS neurons. For this, we treated 10 days in vitro (DIV) embryonic rat cortical neuron cultures with scrambled (E7P7) or G3BP1 192-205 (EP_7_) CPP for 1 h and then assayed the effect of the CPPs on G3BP1 and other stress granule proteins by confocal microscopy. The G3BP1 192-205 (EP_7_) CPP significantly reduces both the density and size of axonal G3BP1 granules (**Fig. 5A-C**). G3BP2 and FMRP axonal granule density were decreased by the 192-205 (EP_7_) CPP, but granule size was not affected (**Fig. 5D-F, Suppl. Fig. S6A-C**). Interestingly, the G3BP1 192-205 (EP_7_) CPP did not show any significant effect on axonal FXR1 granule density or size (**Suppl. Fig. S6D-F**). Overall, these data indicate that the G3BP1 CPP promotes the disassembly of axonal LLPS for multiple stress granule-linked proteins.

**Figure 5:**
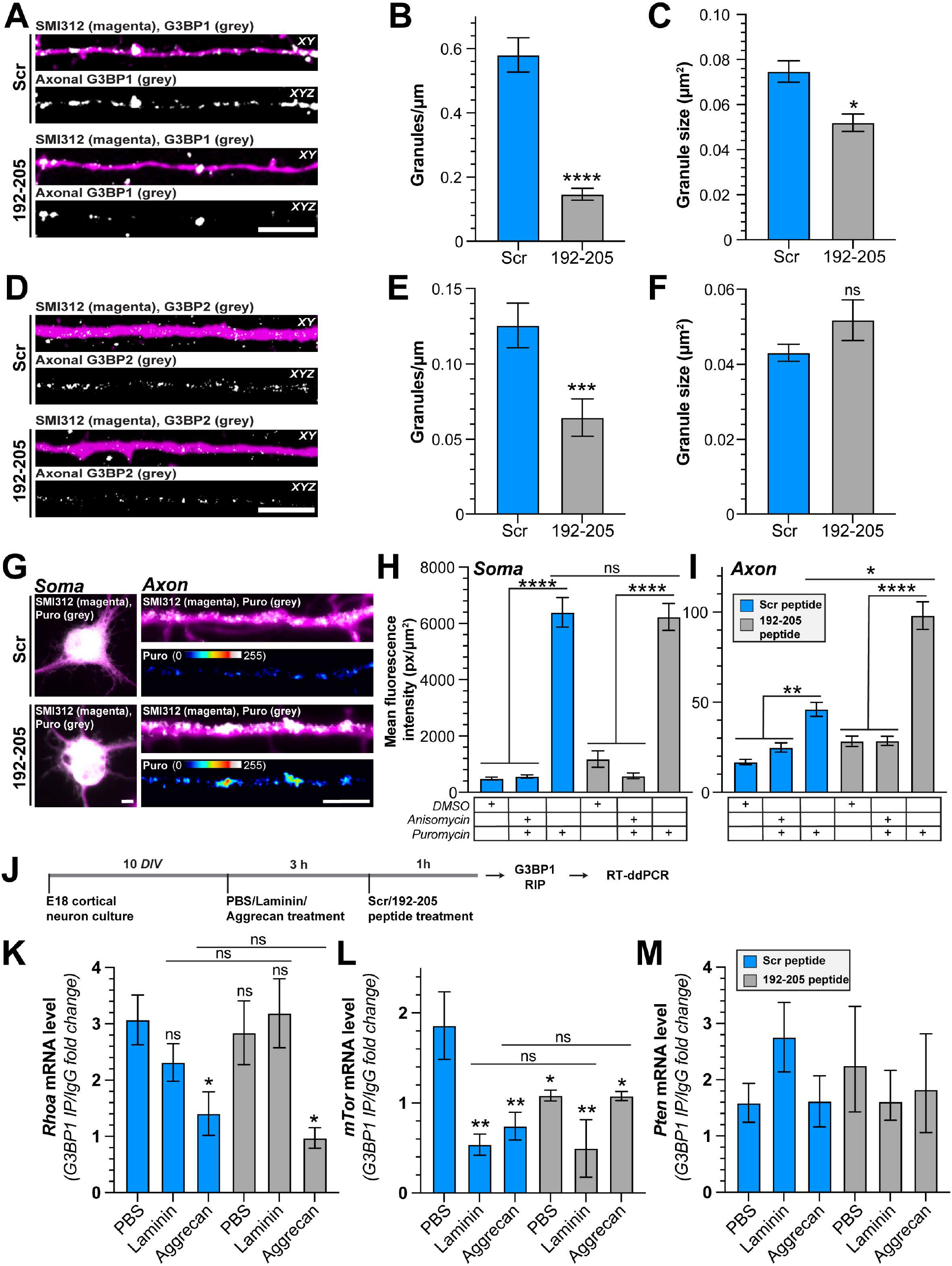
G3BP1 granule disassembling CPP affects other stress granule proteins, activates axonal protein synthesis, and shows specificity for mRNA release. ***A-C***, Representative exposure-matched confocal images of axons of embryonic rat cortical neurons labeled with axonal neurofilament marker SMI312 and G3BP1 are shown (A). The upper panel of each image pair shows merged signals in a single XY plane, and the lower panel shows the XYZ projection of G3BP1 signals that overlap with SMI312 across individual Z sections. Quantitation of the density (B) and size (C) of axonal G3BP1 granules shown as mean ± SEM (C) (N ≥ 33 axons across three biological replicates, ****p ≤ 0.0001 by Mann Whitney U test in B and N ≥ 79 granules across three biological replicates, *p ≤ 0.05 by Mann Whitney U test in C) [scale bar = 5 μm]. ***D-F***, Representative exposure-matched confocal images of axons of embryonic rat cortical neurons labeled with axonal neurofilament marker SMI312 and G3BP2 (D) are shown as in A. Quantitation of the density (E) and size (F) of axonal G3BP2 granules shown as mean ± SEM (N ≥ 38 axons across three biological replicates, ***p ≤ 0.001 by Mann Whitney U test in E and N ≥ 76 granules across three biological replicates, p values by Mann Whitney U test in F). For FMRP and FXR1 granule size after CPP treatment, see Supplemental Figures S6A-F [scale bar = 5 μm]. ***G-I***, Exposure-matched epifluorescence images for SMI312 + Puromycin immunostained embryonic rat cortical neurons after CPP treatment are shown (G). Quantitation of the amount of puromycin in soma (**H**) and axons (**I**) of embryonic rat cortical neurons after CPP treatment shown as mean ± SEM (N ≥ 28 neurons across three biological replicates; ****p ≤ 0.0001, **p ≤ 0.01, *p ≤ 0.05 Kruskal Wallis test with Dunn’s multiple comparisons). For puromycin quantitation in Schwann cells after CPP treatment, see Supplemental Figures S6G, H [scale bar = 5 μm]. **J-M**, Schematic of experimental paradigm for analyses of RNAs coprecipitating with G3BP1 (**J**). Quantitation of fold change in mRNA levels of Rhoa (**K**), mTor (**L**), and Pten (**M**) interacting with G3BP1 after bath-application of laminin or aggrecan ± CPP treatment shown as mean ± SEM (N ≥ 3 animals, *p ≤ 0.05, **p ≤ 0.01 by Ordinary One-way ANOVA with Dunnett’s multiple comparisons).

We previously showed that the 190-208 G3BP1 CPP increases axonal but not cell body protein synthesis in adult DRG neurons (18). Thus, we asked if the G3BP1 192-205 (EP_7_) CPP can modulate protein synthesis in CNS neurons. 30 min exposure to the G3BP1 192-205 (EP_7_) CPP specifically increases axonal protein synthesis in 10 DIV embryonic rat cortical neuron cultures without any significant effect on soma protein synthesis (**Fig. 5G-I**). Because this peptide also penetrates glial cells, we asked if protein synthesis in Schwann cells of mixed adult DRG cultures would be affected by G3BP1 192-205 (EP_7_) CPP treatment. There was no significant change in protein synthesis in the Schwann cells by puromycinylation comparing 192-205 (EP_7_) vs. scrambled (E_7_P_7_) CPP-treated cultures (**Suppl. Fig. S6G-H**). Taken together, these data suggest that the G3BP1 granule disassembling CPP specifically increases protein synthesis in the axonal compartment.

Considering the effect of RhoA inhibition shown in Figures 4 and S5, we asked if the G3BP1 granules in cortical neurons contain *RhoA* mRNA and if this interaction is affected by growth-inhibiting vs. growth-promoting stimuli. Analysis of RNA coimmunoprecipitating with G3BP1 (i.e., RNA co-immunoprecipitation or RIP) by reverse-transcriptase droplet-digital PCR (RT-ddPCR) data shows that treatment of cortical neurons with aggrecan significantly reduces the interaction of *Rhoa* mRNA with G3BP1 and the G3BP1 192-205 (EP_7_) CPP has no apparent effect on G3BP1-*Rhoa* interaction (**Fig. 5K, Suppl. Fig. S6I**). Considering the known role of the mTor pathway in modulating CNS and PNS axon growth (3-9), we asked if *mTor* and *Pten* mRNAs might show G3BP1 association that is altered by growth-inhibiting vs. growth-promoting stimuli. Treatment with laminin, aggrecan, or the G3BP1 192-205 (EP_7_) CPP attenuated G3BP1-*mTor* mRNA interaction but had no significant effect on G3BP1-*Pten* mRNA interaction (**Fig. 5L, M**). Taken together, these data emphasize that the specificity of response to G3BP1 granule disassembly is modulated by the neuronal growth environment, providing a likely explanation for the differences in axon growth promotion seen with granule disassembly in CNS vs. PNS extracellular environments.

## DISCUSSION

In contrast to the PNS, low capacity for axon growth after injury combined with a growth-inhibitory extracellular milieu in the injured brain and spinal cord limits the mature CNS neurons’ axon regeneration after traumatic CNS injuries (2). Consequently, interventions are needed to coax CNS axons to regenerate. PNS axons can spontaneously regenerate but grow very slowly at 1-4 mm per day (1). We previously showed that decreasing LLPS of RNA-protein granules containing the core stress granule protein G3BP1 accelerates the regeneration of injured PNS axons (18, 20). Here, we show that expression of G3BP1’s conserved B-domain (amino acids 141-220) accelerates regeneration of reticulospinal axons in the growth-permissive environment of a peripheral nerve grafted into the hemisected spinal cord and is sufficient to enable injured RGCs to regenerate axons in the growth-inhibitory CNS environment of the injured optic nerve. We previously found that this B-domain functions as a dominant negative agent by preventing axonal G3BP1 granule assembly and increasing axonal protein synthesis (18). Since G3BP1 B-domain expression in reticulospinal neurons and RGCs decreased granule formation by endogenous G3BP1 in their axons, G3BP1 B-domain expression in the CNS neurons likely increases regeneration by releasing axonal mRNAs encoding regeneration-associated proteins that are normally sequestered in G3BP1 granules. Notably, depletion of the *C. elegans* orthologue of TIA1, TIAR-2, similarly increased regeneration following axotomy by decreasing RNA-protein granules and increasing axonal protein synthesis (41). Together with the findings herein, this observation emphasizes that mRNA sequestration impedes mammalian CNS axon regeneration just as previously shown for mammalian PNS nerve and for invertebrate axon regeneration (18, 41).

Some interventions that increase PNS axon regeneration have proven successful for promoting CNS axon regrowth, though this is often limited in terms of the extent of growth promotion. For example, conditioning lumbosacral sensory neurons by sciatic nerve crush injury accelerates axon regeneration after subsequent PNS injury and brings a degree of axon regrowth after dorsal column lesion in the spinal cord (42). This phenomenon is attributable to the entry of inflammatory cells into the injured PNS, which elevates levels of cell-derived growth factors including oncomodulin and SDF-1 (43). The PTEN→mTOR pathway has been manipulated to increase CNS and PNS axon regeneration either by attenuating PTEN activity or downstream interventions that more directly increase mTOR activity (4, 7). PTEN activity can modify gene transcription (44), however, the increase in mTOR activation with PTEN deletion also converges on the translation machinery to globally increase cap-dependent protein synthesis (45). The resulting activation of the mTOR pathway can promote axon regeneration even in injured corticospinal tract axons that normally show very poor growth (8). *mTor* mRNA is locally translated in PNS axons in response to an injury-induced increase in axoplasmic Ca^2+^ (20, 46). This increase in axonal mTOR protein supports neuronal survival after injury (46) but is also needed for subsequent translation of axonal *Csnk2a1* mRNA, with newly generated CK2α phosphorylating G3BP1 and triggering G3BP1 granule disassembly to increase axonal protein synthesis (20). *mTor* mRNA is also locally translated in axons of cultured cortical neurons (47) and the mRNA and protein are present in growth cones of developing CNS neurons *in vivo* (48). We find that *mTor* mRNA coprecipitates with neuronal G3BP1 protein and exposure to either aggrecan or laminin depletes the mRNA from G3BP1 interaction. *mTor* mRNA was also released from G3BP1 interaction by the G3BP1 granule disassembling CPP. In contrast to PNS and other CNS neurons, a large proportion of RGCs die after optic nerve injury; similar to what we show with G3BP1 B-domain expression in RGCs, deletion of the murine *PTEN* gene was shown to increase survival of axotomized RGCs in addition to promoting regeneration (9). Thus, aggrecan-induced release of *mTor* mRNA from G3BP1 interaction upon G3BP1 granule disassembly may indeed prevent the death of injured RGCs that were transduced with the AAV2-G3BP1 B-domain.

Though we have previously shown that G3BP1 B-domain and 190-208 CPP decrease axonal G3BP1 granules and increase translation of protein products that promote axon growth (18), it is not yet clear how either disrupts the G3BP1 granules. The B-domain is one of G3BP1’s three intrinsically disordered regions (IDR) and is required for LLPS by G3BP1 (49). This region is well-conserved across vertebrate species, with mouse showing more than 85% homology to human, 67% to xenopus, and 46% to zebrafish based on G3BP1 sequences available in UniProt (50). The rodent 190-208 and human 191-209 sequences lie within G3BP1’s IDR1, which intramolecularly interacts with the protein’s IDR3 (49, 51). CPPs have been employed to disrupt protein-protein interactions (52). For example, CPPs were used to disrupt mutant Huntingtin’s (HTT) protein-protein interactions in a Huntington disease model and DJ-1’s (PARK7) interactions in a Parkinson disease model (53, 54). Though the G3BP1 B-domain (IDR1) was shown to intramolecularly interact with the protein’s IDR3 (49, 51), we do not detect coprecipitation of G3BP1 with a biotinylated 190-208 G3BP1 CPP, even when testing recombinant G3BP1 combined with crosslinking (raw data will be submitted to https://odc-sci.org). Thus, interruption of G3BP1’s intramolecular interactions or interactions of multiple G3BP1 proteins with one another is an unlikely mechanism. G3BP1 is a core stress granule protein that we previously showed interacts with other stress granule proteins in PNS sensory axons including TIA1 (18). We show that core functional amino acids of the G3BP1 CPP (i.e., the EP7 repeat region, rat amino acids 192-205) disrupts G3BP1, G3BP2, and FMRP, but not FXR1 granules, in axons of cultured cortical neurons. Similar to the CPP not binding to G3BP1, we also do not see coprecipitation of G3BP2 with the 190-208 CPP (raw data will be submitted to https://odc-sci.org); thus, future studies will be needed to determine how the 190-208 CPP disassembles G3BP1 granules. Depletion and knockout approaches have defined several core stress granule proteins and proximity biotinylation approaches have uncovered many G3BP1 interacting proteins (49, 51, 55-57). Axonal G3BP1 colocalizes with TDP43, and rodent 190-208 G3BP1 CPP triggers disassembly of pathological TDP43 aggregates in axons of iPSC-derived human motor neurons (58). Additionally, G3BP1 binds to the adapter protein Annexin A11, which allows the RNP complex to be transported along axons by hitchhiking on lysosomes/endosomes as a means for delivering mRNAs into axons (59). Since we see that sustained exposure to the 190-208 G3BP1 CPP facilitates axon growth and delivery of new mRNAs into axons that are needed for regeneration (10), the 190-208 CPP likely does not interrupt G3BP1-AnnexinA11 interactions. Additional studies beyond the structure-activity relationship analyses included here will be needed to uncover how the G3BP1 CPP and B-domain disrupt G3BP1 granules.

The rodent 190-208 G3BP1 CPP contains seven Glu-Pro repeats, and a CPP with just the seven Glu-Pro repeats (rodent G3BP1 amino acids 192-205) is sufficient for neurite growth-promoting activity. Acidic residues with alternating prolines are essential for this growth promotion, because a CPP with Asp residues replacing Glu has full activity, but its activity is lost on scrambling the Glu-Pro repeats. Tertiary structures of rodent 190-208 and human 191-209 G3BP1 sequences generated with *Pepfold3* (60-62) predict a kink in the peptide backbone at each Pro with the acidic side chains of each acidic residue projecting peripherally from the backbone (**Suppl. Fig. S3Ei-ii**), which is lost in the scrambled rodent 190-208 G3BP1 CPP (**Suppl Fig. S3Eiii**). These predicted 3D structure differences of the rodent 190-208 and human 191-209 G3BP1 CPPs compared to scrambled rodent 190-208 CPP, as well as the rodent 147-168^S149E^ G3BP1 CPP (**Suppl. Fig. S3Eiv**), may contribute to differences in their growth promoting abilities.

It is intriguing that the 190-208 peptide increased branching of axons in the PNGs when presented in the non-permissive environment of the injured spinal cord (**Fig. 3A-D**) and for primary neurons cultured on non-permissive CSPG substrate compared to elongating growth on permissive laminin substrate (**Fig. 4B, Suppl. Fig S5A-D**). This effect is in contrast to G3BP1 B-domain expression that must begin before injury – this may point to different mechanisms underlying prevention of G3BP1 granule formation (i.e., viral-based B-domain expression before injury) vs. disassembly of formed G3BP1 granules (i.e., 190-208 G3BP1 CPP administration after injury). Interestingly, inhibition of ROCK partially reversed the increased branching seen in the 190-208 G3BP1 CPP-treated neurons grown on aggrecan (**Fig. 4C-D, Suppl. Fig S5E**). Activation of RhoA causes actin filament disassembly (63), and *RhoA* mRNA is locally translated in axons (64) with previous studies showing that CSPGs stimulate axonal *RhoA* mRNA translation in axons (31). This raises the possibility that the axon’s environment determines either the complement of mRNAs released from G3BP1 granules or which released mRNAs are translated in the axons. *RhoA* mRNA has been detected in G3BP1-RNA coimmunoprecipitations (65). Axon branching requires local reorganization of the cytoskeleton (66), and increasing RhoA protein locally could support by increase axon branching by disassembling actin microfilaments. CSPG stimulation can increase axoplasmic Ca^2+^ (67, 68) and axoplasmic Ca^2+^ levels can determine which axonal RNAs released from G3BP1 granules are translated through the phosphorylation of the translation factor eIF2α in axons (20). Consistent with this, we find that *RhoA* mRNA coprecipitates with neuronal G3BP1 protein and this interaction is decreased in the presence of aggrecan but was not affected by the G3BP1 granule disassembling CPP. Interestingly, RhoA activation was shown to increase formation of TIA1- and FMRP-containing stress granules in other cellular systems (69), and CSPG-stimulated RhoA can activate the cytoplasmic histone deacetylase HDAC6 in distal axons (68). Although Y27632 has been extensively used in the past to inhibit ROCK, it is now known to inhibit multiple other kinases to some extent (70). So, we cannot exclude the off-target effects of Y27632 contributing to the rescue in axon growth and branching that we see in Fig. 4C-D. Deacetylation of G3BP1 lysine 376 by HDAC6 promotes G3BP1 granule assembly (71). Considering that the rat 190-208 G3BP1 and human 191-209 sequences do not include the post-translationally modified Ser 149 or Lys 376, we suspect that these CPPs do not alter post-translational modifications of G3BP1. Taken together, our data raise the possibility that successful long-distance CNS axon regeneration with G3BP1 granule inhibition will require simultaneous inhibition of RhoA.

In summary, our findings show that G3BP1 granules contribute to failed axon regeneration after optic nerve and spinal cord injury. Thus, disassembly of G3BP1 granules, as we have done here using G3BP1 B-domain expression or the 190-208 G3BP1 CPP, could bring a new strategy to promote neural repair following brain and spinal cord injuries, either on their own or strategically combined with other interventions. Importantly, the G3BP1 190-208 CPP strategy used here is effective when delivered 2 days after PNS injury and at least 30 days after spinal cord injury, which elevates the potential clinical relevance of these reagents for neural repair treatments. Structure-activity relationship studies point to alternating acidic residue (Glu or Asp) and Pro repeats as the functional component of the 190-208 G3BP1 CPP. This presents opportunities for further refinement of the CPP activity, and since the 191-209 human G3BP1 CPP promotes axon growth in human iPSC-derived CNS neurons, we anticipate that similar reagents may have high translational value for neural repair in human patients.

## MATERIALS AND METHODS

### Animal use

Institutional Animal Care and Use Committees of University of South Carolina, Rutgers University, Boston Children’s Hospital, Emory University, University of California San Diego, Texas A&M University, and Drexel University approved all animal procedures. Sprague Dawley rats (SD; 175-250 g) were used for sciatic nerve injury, and spinal cord injury, and wild type 129s mice were used for optic nerve injury experiments. Survival surgery methods are described below.

### Peripheral nerve injury

For crush injury of the sciatic nerve, 5% isoflurane in 1 L/min oxygen was used for anesthesia and maintained at 2% isoflurane in 1 L/min oxygen. Anesthetized rats or mice were subjected to a manual crush at mid-thigh two times at 15 sec each using fine jeweler’s forceps with a 30 sec interval, and sciatic nerves were isolated 1 week later as previously described (72).

### Analysis of muscle reinnervation after peripheral nerve injury

Following nerve injury and CPP injections, the extent of reinnervation of the hindlimb muscles was evaluated using in vivo electromyography (EMG). Anesthesia was induced with 5% isoflurane in 1 L/min oxygen and maintained at 2% isoflurane in 1 L/min oxygen.

### Spinal cord injury stem cell graft

Adult female F344 Fischer rats received C4 dorsal column lesion spinal cord injury. Surgeries were conducted under deep anesthesia using a combination of ketamine (50 mg/kg), xylazine (2.6 mg/kg), and acepromazine (0.5 mg/kg). After performing a laminectomy, a tungsten wire knife (McHugh Milieux, Downers Grove, IL) was positioned 0.6 mm lateral to the midline and inserted to a depth of 1 mm beneath the dorsal surface of the spinal cord. The knife’s arc was then extended 1.5 mm and elevated to transect the dorsal columns as previously described (73). This injury model transects over 95% of corticospinal axons (74).

### Analysis of reticulospinal tract axon regeneration in PNGs

For testing expression of G3BP1 B-domain in reticulospinal neurons, AAV5-G3BP1-BFP or AAV5-B domain-BFP was microinjected into the brainstem of female SD (225 g, obtained from Charles River) on day 1 of experimentation. Control rats received AAV5-GFP. Rats received three 1 µl injections at 1 mm rostral-caudal separations into medial-ventral gigantocellular areas of the reticular formation (see Figure 1B).

### Optic nerve injury and RGC survival analysis

Optic nerve regeneration and RGC survival were investigated in 129s mice. The optic nerve was exposed and crushed 1-2 mm behind the orbit using jeweler’s forceps using methods that have been previously described (26, 75-77).

### Cell culture

For primary neuronal cultures, DRGs were harvested in Hibernate-A medium (BrainBits; Cordova, TN) and then dissociated as described (72). After centrifugation and washing in DMEM/F12 (Life Technologies), dissociated ganglia were resuspended in DMEM/F12, 1 x N1 supplement (Sigma), 10% fetal bovine serum (Hyclone; Logan, UT), and 10 μM cytosine arabinoside (Sigma). Dissociated DRGs were plated immediately on poly-L-lysine (Sigma) + laminin (Sigma-Millipore, St. Louis, MO) or poly-L-lysine + aggrecan (R&D Systems, Minneapolis, MN) coated surfaces (20, 72). For Schwann cells, dissociated DRGs were cultured as above without cytosine arabinoside. Schwann cells were identified at 2 DIV by co-immunostaining for GFAP in the puromycinylation studies. The use of human iPSCs was approved by the Institutional Review Board at the University of South Carolina. Human iPSCs from apparently healthy euploid individuals were obtained from Dr. Alberto Costa (Case Western Reserve University; Cleveland, Ohio) (78).

### Viral expression constructs

AAV2 and AAV5 preparations were generated in Vigene Biosciences/Charles Rivers Labs (Wilmington, MA) and the viruses were titrated in DRG cultures by incubating with 1.8-2.8 × 10^10^ particles of AAV2/5 overnight. Generation of the viral constructs has been described previously (18).

### Generation of Tat-tagged G3BP1 B domain peptides

Eight peptides were generated from the rat G3BP1 B domain sequence (amino acids 140-220; UniProt ID # D3ZYS7_RAT) by Peptide 2.0 (Chantilly, VA) and Bachem Americas, Inc. (Torrance, CA).

### Puromycinylation assays

At 2 DIV, Schwann cells were starved using DMEM/F12 and 1x L-Glutamine for 3 h. 10 µM Scrambled or 192-205-CPPs were added 15 min before completion of starvation. Schwann cells were stimulated with 10% FBS and 1x N1 supplement. For translation inhibition controls, anisomycin (Sigma, 40 µM) or DMSO was added 30 min before completion of starvation for a total duration of 45 min. Puromycin (Sigma, 2 µg/ml) was added at the time of stimulation for a total duration of 15 min (79, 80). Cultures were then fixed with 4% PFA, and immunofluorescence for GFAP and puromycin was performed.

### RNA immunoprecipitation (RIP)

10 DIV cultured rat cortical neurons were treated with PBS, laminin (5 µg/ml), or aggrecan (5 µg/ml) in solution for 4 h were used for RIP studies. Neurons were collected and washed twice with cold PBS and were lysed in RIP buffer (containing 100⍰mM KCl, 5⍰mM MgCl2, 10⍰mM HEPES [pH 7.0], 1⍰mM DTT, and 0.5% NP-40 supplemented with 1⍰× protease inhibitor cocktail (Roche) and RNasin Plus (Invitrogen), by pipetting up-down and incubating on ice for 10 min. Lysate was cleared by centrifugation at 12,000×g for 20⍰min. Pre-washed Protein A/G Dynabeads (ThermoFisher) were incubated at 4°C for 1 h with anti-rabbit IgG (Jackson Immunoresearch) or anti-G3BP1 antibody (Fisher Scientific) at 0.67 µg/ml each in RNAse-free PBS and then washed with RIP buffer three times. The antibody-bound beads were resuspended with 500 μl of NT2 buffer (containing 50 mM Tris-HCl [pH 7.4], 150⍰mM NaCl, 1 mM MgCl2 and 0.5% NP-40 supplemented with 1⍰× protease inhibitor cocktail and RNasin Plus (Invitrogen). 10% of cleared lysate was kept as input, and the rest was divided in half for control IgG RIP or G3BP1 RIP. Lysates were added to antibody-protein A/G beads complex in NT2 buffer and incubated at 4°C for 4 h. The immunocomplexes were washed with NT2 buffer 6 times (18). The bound RNAs were purified and analyzed by RTddPCR (*see below*).

### RNA isolation and RT-ddPCR analysis

RNA was isolated from inputs and immunoprecipitates using Trizol (Invitrogen) and chloroform (MP biomedicals) and subsequently precipitated using isopropanol and washed with 75% ethanol. Reverse transcription was done using LunaScript® RT SuperMix Kit (NEB). ddPCR products were detected using QX200™ ddPCR™ EvaGreen Supermix (Biorad) and QX200® droplet reader (Biorad).

Custom transcript-specific primer sets used for ddPCR were as follows: *RhoA*, forward primer - CCAGACTGGACTGAGGAAATAG and reverse primer - GAAAGGAGCACTGTGACTTAGA; *mTor*, forward primer - TGTCTGATTCTTACCACGCA and reverse primer - CTCTTTGGCCAGGGTCTCAT; and *Pten*, forward primer - AGCGTGCGGATAATGACAAG and reverse primer - GGATTTGATGGTTCCTCTACTG.

### Immunoblotting

For immunoblotting, proteins were transferred from gels to nitrocellulose membranes and blocked for 1 hour with 5% milk in Tris-buffered saline with 0.1% Tween 20 (TBST) at room temperature. Membranes were incubated overnight at 4°C with Rabbit anti-G3BP1 antibody (1:2,000; Sigma) diluted in the blocking buffer while rocking. After washing with TBST, membranes were incubated for 1 hour at room temperature with HRP-conjugated anti-rabbit IgG secondary antibody (1:5,000; Jackson ImmunoResearch). Following washes with TBST, immunocomplexes were visualized using Clarity™ Western ECL Substrate (Biorad). Immunofluorescent staining – All procedures were performed at room temperature (RT) unless specified otherwise. Cultured neurons were fixed in 4 % paraformaldehyde (PFA) in phosphate-buffered saline (PBS), permeabilized in 0.3% TritonX, and incubated with primary and secondary antibody solutions in blocking buffer as described previously (81).

### Image analyses and processing

For analyses of stress granule protein levels in tissues, z planes of the xyz scans of spinal cord sections were analyzed using *ImageJ*. Axons, neurons, and Schwann cells were identified using specific markers, and SG protein colocalization with neuronal or glial cell makers was assayed as described previously (14).

### Statistical analyses

One-way ANOVA, Two-way ANOVA, or an equivalent non-parametric test was used to compare means of > 2 independent groups and Student’s *t*-test or Mann-Whitney test was used to compare between 2 groups. p values of ≤ 0.05 were considered as statistically significant. Z scores were calculated using the formula 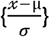, where *x* = sample mean, *µ* = population mean, and σ = population standard deviation.

### Other Methods

Detailed methods can be found in the *SI Appendix, Supplemental Materials and Methods*.

## Acknowledgements

The authors thank members of the K. Welshhans, F. Poulain and D.S. Smith laboratories at the University of South Carolina for constructive discussions. JL Twiss is the incumbent SmartState Chair in Childhood Neurotherapeutics at the University of South Carolina.

## Data and materials availability

The data that support the findings of this study will be deposited in a public repository (https://odc-sci.org). PKS and JLT hold a US patent on use of the G3BP1 cell permeable peptide for axon regeneration and neurodegeneration. So, request for this reagent will be available by materials transfer agreements (MTAs) with the University of South Carolina.

