## Supplemental text for "Disruption of G3BP1 Granules Promotes Mammalian CNS and PNS Axon Regeneration"

### Main Manuscript for

**This PDF file includes:** Supporting Text
Suppl Figures S1 to S6
SI References

### Supporting Information Text

#### Supplemental Materials and Methods.

***Peripheral nerve injury*** **–** For *in vivo* cell permeable peptide (CPP) injections, 1 µl of 1 mM cell permeable peptides (diluted in 10 µl phosphate-buffered saline; PBS) were injected into sciatic nerves proximal to the crush site 2 days after injury. Concentration for *in vivo* peptide delivery into the nerve was determined based on estimated volume of a 1.5 mm nerve cylinder and diffusion of 2.5 mm in each direction giving a nerve volume of 8.84 $mm^3$. 10 µl solutions from above were injected in 3 different sites (~3.3 µl in each site) within 500 µm proximal to the injury site using Hamilton syringes to approximate an in vivo peptide concentration of 113 µM. The needle tip was inserted horizontally for 2-3 mm into the sciatic nerve for peptide injection over a 1-2 min period.

***Analysis of muscle reinnervation after peripheral nerve injury*** **–** Bipolar fine wire EMG electrodes were constructed from insulated nichrome wire (A-M Systems, Sequim, WA). The insulation over the distal 1 mm of the tips was removed by scraping with a scalpel blade and the tips of the two wires were staggered by 1 mm. Electrodes were then placed into the mid-belly of the lateral head of the gastrocnemius (LG) and tibialis anterior (TA) muscles using a 25 Ga hypodermic needle. Once in place, the needle was removed, and the wires were connected to the differential amplifiers. To stimulate the sciatic nerve, a small skin incision was made just inferior to the ischial tuberosity, exposing the sciatic nerve as it passes between the gluteal and hamstring muscles and proximal to the crush injury site. Two unipolar needle electrodes (Neuroline monopolar, 28 Ga; Ambu/AS, Columbia, MD) were placed on either side of the sciatic nerve, separated from each other by approximately 1 mm. Lead wires from the needles were connected to an optically isolated constant voltage stimulator under computer control (1).

Evoked EMG activity from LG and TA was then recorded during sciatic nerve stimulation. Stimulation and recording were controlled by a laboratory computer system running custom software written in Labview® (Emerson, St. Louis, MO). Ongoing EMG activity in the LG was sampled at 10 kHz; when the rectified and integrated voltage over a 20 msec period fell within a user-defined range, a 0.3 msec duration stimulus pulse was delivered to the nerve via the needle electrodes. Muscle activity was sampled and recorded from 20 msec prior to the stimulus until 100 msec after the stimulus. Stimuli were delivered no more frequently than once every 3 sec to avoid muscle fatigue. A range of stimulus intensities was applied in each experiment to sample evoked muscle activity from sub-threshold to supramaximal. In a typical experiment, approximately 200 stimulus presentations were studied. At the end of each experiment, all electrodes were removed, and the skin was incision closed with sutures.

The recorded compound muscle action potentials (CMAPs) in LG and TA evoked by sciatic nerve stimulation were analyzed off-line. The amplitude of the evoked M waves was measured as the average rectified voltage within a defined time window after the stimulus application. In intact anesthetized animals, this window is 0.5-2 msec. After nerve crush, M waves evoked from sciatic nerve stimulation are, by definition, generated by reinnervated muscle fibers. The latency and duration of these potentials are longer than those found in intact animals (2), and thus, the time window used to measure the amplitude of the M waves was adjusted accordingly. Recordings were made from intact animals, and 2, 4, 6, and 8 wk after nerve crush. At each time point, the amplitude of the largest evoked CMAP was determined and scaled to CMAPs recorded from that animal prior to nerve crush. Means of these scaled responses from animals in which nerves were treated with CPP 168-189 were compared to animals in which nerves were treated with CPP 190-208 at each time studied.

***Spinal cord injury stem cell graft* –** Immediately following injury, rats received transplantation of dissociated neural progenitor cells obtained from the spinal cord (caudalized) or telencephalon (rostralized) of E14 GFP+ F344 rat embryos. Neural progenitor cells were isolated from embryonic tissue by incubation with 0.125% trypsin, gentle mechanical dissociation, and filtering through a 40 µm cell filter as previously described (3, 4). A total of $10^6$ cells in HBSS were transplanted into the lesion site. Four weeks after SCI and transplantation, rats received cortical injections of 10% biotinylated dextran amine (BDA) across 16 sites into bilateral primary forelimb motor cortices. Animals were sacrificed four weeks later.

***Analysis of reticulospinal tract axon regeneration in PNGs* –** On day 8 the tibial branch of the sciatic nerve of donor SD rats was ligated and cut just distal to its branch point from the sciatic nerve. Seven days later a 10 mm length of degenerated tibial nerve was removed from the donor rat and the distal end was inserted into a C4 hemisection lesion cavity that was prepared by aspiration in a recipient rat. The graft was secured by suturing the perineurium to the dura mater. The distal end of the PNG was left unapposed, lying on top of the adjacent, distal vertebral processes. On day 35 of experimentation (20 days after SCI-PNG) rats were euthanized (see below for details).

For CPP delivery to injured spinal cord axons proximal to PNG, 1 µl of 1 mM 168-189 peptide or 190-208 peptide was microinjected into the spinal cord 1 mm rostral to a fresh C4 hemisection lesion cavity (see Figure 3A). This was estimated to provide a concentration of 28 µM in the tissue. Approximately 20 min later one end of a predegenerated tibial nerve from a donor rat was inserted into the lesion cavity, apposed to the proximal cavity wall and secured by suturing perineurium to the dura mater. The distal end of the PNG was left unapposed, lying on top of the adjacent, distal vertebral processes. On day 18 of experimentation (10 days after SCI-PNG) rats were euthanized (see below for details).

For CPP delivery to regenerating axons within PNGs, 1 µl of 1 mM 190-208 peptide or scrambled peptide was microinjected into the PNG through a fine glass cannula attached to a Hamilton syringe into the unapposed distal end of the PNG approximately 30 days after the PNG had been apposed to the rostral wall of the injured C4 spinal cord (see Figure 3E). This was estimated to provide a concentration of 113 µM in the tissue. The cannula tip was inserted horizontally for 2-3 mm into the PNG for peptide injection over a 10-minute period. After withdrawing the cannula, the PNG was moved aside and a 1 $mm^3$ dorsal quadrant cavity at the C5-6 interface (between entry zones of dorsal root axons) was created by aspiration. The distal end of the PNG was trimmed by 1 mm and inserted into the fresh cavity and secured by suturing perineurium to dura mater. Three weeks after apposing the distal end of the graft rats were euthanized (see below for details).

For all experiments rats received sustained-release Buprenorphine (1 mg/kg, subcutaneously [SC], Zoopharm, Laramie, WY) for analgesia and the antibiotic Cefazolin (160 mg/kg SC 2x daily for 5 days; Sandoz, Princeton, NJ). Two days prior to placement of PNGs rats began a regimen of daily Cyclosporine A injections (10 mg/kg SC; Teva Czech Industries, Sellersville, PA) to prevent graft rejection. At the end of each experiment rats were euthanized with an overdose of Euthasol (390 mg/kg sodium pentobarbital and 50 mg/kg phenytoin, intraperitoneal [IP]), perfused with physiological saline followed by 4% paraformaldehyde. The spinal cord with attached PNG was removed and transferred to 30% sucrose in 0.1 M phosphate buffer at 4°C before preparing cryostat sections and performing immunohistochemical staining and microscopic analysis.

***Optic nerve injury and RGC survival analysis*** **–** In brief, the optic nerve was exposed and crushed 1-2 mm behind the orbit using jeweler's forceps. In cases involving gene therapy, the gene of interest was packaged into adeno-associated virus (AAV2) at a titer of $10^{12}$-$10^{13}$ pfu/ml and injected in a volume of 3 µl intraocularly 2 weeks prior to optic nerve crush (see Figure 1E) to allow time for virally encoded constructs to be expressed within RGCs at adequate levels. Care was taken to avoid injuring the lens, which can profoundly alter levels of axon regeneration (5).

After a survival period of 2-3 weeks, mice were euthanized, perfused with saline followed by 4% paraformaldehyde, retinas and optic nerves were dissected and postfixed for 1 h. Optic nerves were equilibrated overnight in 30% sucrose, frozen, sectioned longitudinally at 15 µm on a cryostat, and mounted on coated slides.

***Cell culture*** **–** For the CPP effect on cortical neuron cultures, E18 rat cortices were dissected in Hibernate E (BrainBits) and dissociated using the *Neural Tissue Dissociation kit* (Miltenyi Biotec, Auburn, CA). For this, minced cortices were incubated in a pre-warmed enzyme mix at 37°C for 15 min; tissues were then triturated and applied to a 40 µm cell strainer. After washing and centrifugation, neurons were seeded on poly-D-lysine (Sigma) or poly-D-lysine + aggrecan (R&D Systems) coated coverslips. *NbActive-1 medium* (BrainBits) supplemented with 100 U/ml of Penicillin-Streptomycin (Life Technologies), 2 mM L-glutamine (Invitrogen/Thermo-Fisher, Waltham, MA), and 1 X N21 supplement (R&D Systems) was used as culture medium. For puromycinylation, granularity, and RNA immunoprecipitation assays on cortical neuron cultures, E18 rat cortices were dissected in Hank's Balanced Salt Solution (HBSS) (Life Technologies) and dissociated using Trypsin (Life Technologies) as described (6). For this, minced cortices were incubated in pre-warmed trypsin at 37°C for 5 min. After two washes in HBSS, tissues were triturated in Neurobasal with 1x L-Glutamine and B27 supplement (all Life Technologies) and plated on acid-rinsed coverslips (Carolina Biological) pretreated with 100 µg/mL poly-L-lysine (Sigma-Aldrich). After cells adhered to the coverslips (approximately 2 h), the media were changed to fresh Neurobasal with L-Glutamine and B27 supplement. For the puromycinylation and granularity assays, media was supplemented with 1 µg/mL laminin, and depending on the experiment, cells were cultured for 8-11 days at 37°C in 5% $CO_2$.

Human iPSCs were maintained in feeder-free culture conditions on Nunc tissue culture-treated 6-well plates (VWR) coated with Matrigel (Corning, Corning, NY). Cells were maintained in mTeSR1 media (StemCell Technologies, Vancouver, Canada) and passaged every 6-7 days using Gentle Cell Dissociation Reagent (StemCell Technologies).

hiPSC-derived cortical neurons were generated using dual SMAD inhibition in a monolayer layer culture, as previously described (7, 8). This protocol mainly uses two media formulations – Neuronal Maintenance Media (NMM) and Neuronal Induction Media (NIM). NMM was prepared using 1:1 DMEM/F-12 GlutaMAX:Neurobasal (Gibco/Thermo-Fisher, Waltham, MA), 1x N2 supplement (Gibco), 1x B27 supplement (Gibco), 5 µg/mL insulin (Sigma), 1 mM L-glutamine (Gibco), 500 µM sodium pyruvate (Sigma), 100 µM nonessential amino acids solution (Gibco), 100 µM beta-mercaptoethanol (Gibco), 50 U per mL penicillin-streptomycin (Gibco). NIM was prepared by adding 1 µM Dorsomorphin (Tocris, Minneapolis, MN) and 10 µM SB431542 (Reprocell, Beltsville, MD) to NMM. hiPSCs were plated on matrigel-coated wells of a 6-well plate (day 1). The next day (day 0), when cells were 100% confluent, NIM was added for neuronal induction. Cells were maintained in NIM for 12 days with daily feeding. On day 12, which is characterized by the formation of a neuroepithelial sheet, cells were passaged 1:2 using 1 mg/mL dispase into laminin-coated 6-well dishes. The next day, media was replaced with

NMM + 20 ng/mL FGF2, with alternate day feeding. After 4 days of treatment with FGF2, cells were maintained in NMM with alternate day feeding and passaged at a 1:2 split ratio when neuronal rosettes started to meet. On day 25, cultures were dissociated into single cells using accutase and passaged 1:1 onto laminin-coated 6-well dishes. Following this, the cells were passaged every 2-3 days using accutase at a 1:2 split ratio until approximately day 35. For final plating, cells were passaged using accutase. 10,000 cells were plated in NMM on acid-washed coverslips (15 mm; Carolina Biological, Burlington, NC) pre-coated with 100 µg/mL poly-L-lysine (PLL; Sigma) and 10 µg/mL laminin (Invitrogen/Thermo-Fisher).

For peptide treatments, 10 µM CPPs were added to dissociated DRG cultures, and hiPSCs-derived glutamatergic neuron cultures at 16 h after plating. For embryonic cortical cultures, the peptides were added 10 days after plating. Neurite outgrowth was assessed 48 h after addition of peptides.

***Viral expression constructs*** – AAV2 and AAV5 preparations were generated in Vigene Biosciences/Charles Rivers Labs (Wilmington, MA) and the viruses were titrated in DRG cultures by incubating with 1.8-2.8 x $10^{10}$ particles of AAV2/5 overnight.

***Generation of Tat-tagged G3BP1 B domain peptides*** – Peptides were synthesized with N-terminal FITC and N- or C-terminal HIV Tat peptide for cell permeability (9); the Tat sequence was placed at the least conserved end of the sequence based on P-BLAST of vertebrate G3BP1 sequences available in UniProt database. Peptide sequences were (Tat sequence shown in italics): rodent G3BP1 168-189, DDSGTFYDQTVSNDLEEHLEEP-*YGNKKNNQNNN*; rodent G3BP1 190-208, *YGNKKNNNQNNN*-VVEPEPEPEPEPEPEPVSE; rodent G3BP1 190-208$^{E\rightarrow D}$, *YGNKKNNNQNNN*-VVDPDPDPDPDPDPDPVSD; rodent G3BP1 192-205, *YGNKKNNNQNNN*-EPEPEPEPEPEPEP; human G3BP1 191-209, *YGNKKNNNQNNN*-VAEPEPDPEPEPEQEPVSE; Scramble

190-208, *YGNKKNNNQNNN*-VVEEEEEEEPPPPPPPVSE; and rodent G3BP1 147-166$^{S149E}$, EEEEEEVEEPEENQQSPEVV-*YGNKKNNQNNN*.

***Puromycinylation assays*** **–** 10 DIV rat cortical neuron cultures were starved using Neurobasal with 1x L-Glutamine for 3 h (10). 10 µM Scrambled or 192-205-CPPs were added 15 min before completion of starvation. Cortical neurons were stimulated with 1x B27 supplement, 100 ng/µl brain-derived neurotrophic factor (BDNF), and 1 µg/mL laminin. Puromycin was added at the time of stimulation as above for a total duration of 15 min (11, 12). To confirm puromycin signals represented newly synthesized protein, parallel cultures were treated with anisomycin (Sigma, 40 µM) 30 min before completion of starvation. After 15 min puromycin exposure, cells were fixed with 4% PFA and processed for immunofluorescence for puromycin and SMI312.

***RNA isolation and RT-ddPCR analysis*** **–**

***Immunoblotting*** **–**

***Immunofluorescent staining*** **–** Primary antibodies consisted of: RT97 mouse anti-neurofilament (NF; 1:500; Devel. Studies Hybridoma Bank [DHSB], Iowa City, IA), rabbit anti-βIII Tubulin (1:500; Abcam, Boston, MA), Alexa Flour 647-conjugated mouse anti-Puromycin (1:250; Sigma), rabbit anti-G3BP1 (1:100; Sigma), rabbit anti-G3BP2 (1:100; Invitrogen), rabbit anti-FXR1 (1:200; Proteintech), rabbit anti-FMRP (1:50; Cell Signaling), rabbit anti-GFAP (1:200; Bioss), chicken anti-MAP2 (1:3000; Abcam), and SMI312 mouse anti-NF (1:500; Abcam). FITC-conjugated donkey anti-rabbit, Cy3-conjugated donkey anti-mouse, FITC-conjugated donkey anti-mouse, Cy3-conjugated donkey anti-rabbit, Cy5-conjugated donkey anti-chicken, and Alexaflour405-conjugated donkey anti-chicken (all at 1:200; Jackson ImmunoResearch) were used as secondary antibodies.

For quantifying axonal content of G3BP1, and TIA1 in sciatic nerves, peripheral nerve grafts, and optic nerves, samples were fixed for 4 h in 4 % PFA and then cryoprotected overnight in 30 % sucrose, PBS at 4°C. 15 µm cryostat sections were processed for immunostaining as previously described (13). Primary antibodies consisted of rabbit anti-G3BP1 (1:100; Sigma), goat anti-TIA1 (1:100; SantaCruz Bio., Santa Cruz, CA), and RT97 mouse anti-NF (1:300; DSHB). Secondary antibodies were FITC-conjugated donkey anti-rabbit, FITC-conjugated donkey anti-goat, and Cy3-conjugated donkey anti-mouse (both at 1:200; Jackson ImmunoRes.).

For sciatic nerve regeneration analysis, 15 µm thick cryostat sections of sciatic nerve were processed for immunostaining as previously described (13). Primary antibody consisted of rabbit anti-SCG10 (1:200, Abcam). Secondary antibodies were FITC-conjugated donkey anti-rabbit (1:200, Jackson ImmunoRes.).

For reticulospinal tract axon regeneration analysis in PNGs, 15 µm thick cryostat sections were processed for immunostaining. Primary antibody consisted of mouse anti-NF (1:200, DSHB), mouse anti-βIII Tubulin (1:200; Abcam), rabbit anti-SCG10 (1:200; Abcam), rabbit anti-GFP (1:200; Abcam), and rabbit anti-BFP (1:200; Rockland Immuno., Limerick, PA). Secondary antibodies were FITC-conjugated donkey anti-rabbit (1:200; Jackson ImmunoRes.) and Cy3-conjugated donkey anti-mouse (1:200; Jackson ImmunoRes.).

For optic nerve regeneration analysis, 15 µm thick cryostat sections were processed for immunostaining as described previously (14-16). Primary antibody consisted of rabbit anti-BFP (1:200; Rockland) and mouse anti-GAP43 (1:500, Abcam). Secondary antibodies were FITC-conjugated donkey anti-rabbit (1:200; Jackson ImmunoRes.) and Cy3-conjugated donkey anti-mouse (1:200; Jackson ImmunoRes.). For fixed retinas primary antibody consisted of rabbit anti-βIII tubulin (1:100; Abcam), and mouse anti-RBPMS (1:100; Invitrogen). Secondary antibodies were FITC-conjugated donkey anti-rabbit (1:200; Jackson ImmunoRes.) and Cy3-conjugated donkey anti-mouse (1:200; Jackson ImmunoRes.).

All samples were mounted with *Prolong Gold Antifade* or *Prolong Diamond Antifade with DAPI* (Invitrogen/Thermo-Fisher) and analyzed by epifluorescent or confocal microscopy. Leica DMI6000 epifluorescent microscope with ORCA Flash ER CCD camera (Hamamatsu Photonics, Hamamatusu, Japan), Zeiss Axio Observer 7 motorized inverted microscope with Axiocam 820 mono camera, Leica SP8X confocal microscope with HyD detectors, Zeiss LSM980 Confocal Microscope with Airyscan 2 or ImageExpress Micro XLS high content imaging system (Molecular Devices, San Jose, CA) were used for imaging. For quantitation between samples, imaging parameters were matched for exposure, gain, offset and post-processing (2). Thus, the time window used to measure the amplitude of the M waves was adjusted to accommodate this change. Recordings were made from intact animals, and 2, 4, 6, and 8 wk after nerve crush. At each time point, the amplitude of the largest evoked M wave ($M_{max}$) was determined and scaled to $M_{max}$ recorded from that animal prior to nerve crush. Means of these scaled responses recorded from muscles in which motor neurons were treated with the control 168-189 peptide or the 190-208 peptide and were compared at each time studied.

***Image analyses and processing*** – *Colocalization Plug-in* was first used to extract G3BP1 signals that overlap with TIA1 and the extracted colocalizing signal was projected as a separate channel. Further, *colocalization Plug-in* was used for the second time, and the G3BP1-TIA1 colocalization channel was overlapped with the axonal marker (NF) in each plane, to extract the 'axon-only' signal projected as a separate channel (17). For calculating axonal G3BP1 granules in PNGs and optic nerves, XYZ scans from 2 locations along each nerve section were analyzed using *ImageJ.* For calculating axonal G3BP1, G3BP2, FMRP, and FXR1 granules in cortical neurons, only axonal segments ≥ 200 µm from the soma were considered. *ImageJ* segmented line tool was used to mark the axons tracking the NF or SMI312 signal and NF/SMI312 overlapping RBP signals were projected as a separate channel. *Image J* particle analyzer was used to quantify the number and size of the granules. For PNGs, and optic nerves, the axonal G3BP1 granule number and size were averaged for all image locations in each biological replicate.

For puromycinylation quantifications in cortical neurons, soma and the axonal segments ≥ 200 µm from the soma were considered. *ImageJ* polygon tool was used to mark the soma, and the segmented line tool was used to mark the axons tracking the SMI312 signal. For Schwann cells, *ImageJ* polygon tool was used to mark the cells tracking GFAP signal. Mean fluorescence intensity was quantified using *ImageJ* for all puromycinylation assays.

For neurite outgrowth, images from 60 h DRG, or hiPSCs, and 12 days cortical cultures images were analyzed for neurite length and branching using *WIS-Neuromath* (18) with NF, Tubulin βIII, SMI312 immunostained cultures. For reticulospinal axon branching in analyses, high resolution confocal microscopy of the 0 mm PNG sections was performed and further analyzed using Neurolucida 360 and Neurolucida Explorer software (MBF Bioscience, VT). Branching complexity was assessed using the default software parameters.

For optic nerve regeneration analysis, confocal microscopy of anti-GFP/BFP and -GAP43 immunostained optic nerve sections were performed and GAP43-positive regenerating axons were counted manually at 0, 0.5, 1, 1.5 and 2.0 mm from the injury site. 4-5 sections/animal were counted and the number of axons in the entire nerve was estimated using as described (14).

For RGC survival analysis, fixed retinas were immunostained for Tubulin βIII and RBPMS and were divided into four quadrants with radial relieving cuts, mounted on prepared slides, and RGC survival was evaluated by counting the number of RBPMS positive stained cells/mm$^2$ in 8 pre-specified areas of retina (2 areas in each quadrant at 1.0 and 2.0 mm from the center).

To assess regeneration *in vivo*, tile scans of SCG10-stained nerve sections were post-processed by *ImageJ Straighten Plug-in* (http://imagej.nih.gov/ij/). NF positive axon profiles (>30 µm long) were then counted in 30 µm bins at 0.25 mm intervals proceeding distally from the crush site. Crush site was identified by SCG10 staining and DIC imaging. Axon profiles present in the proximal crush site was treated as the baseline, and values from the distal bins were normalized to this to calculate the percentage of regenerating axon profiles.

### SUPPLEMENTAL FIGURES

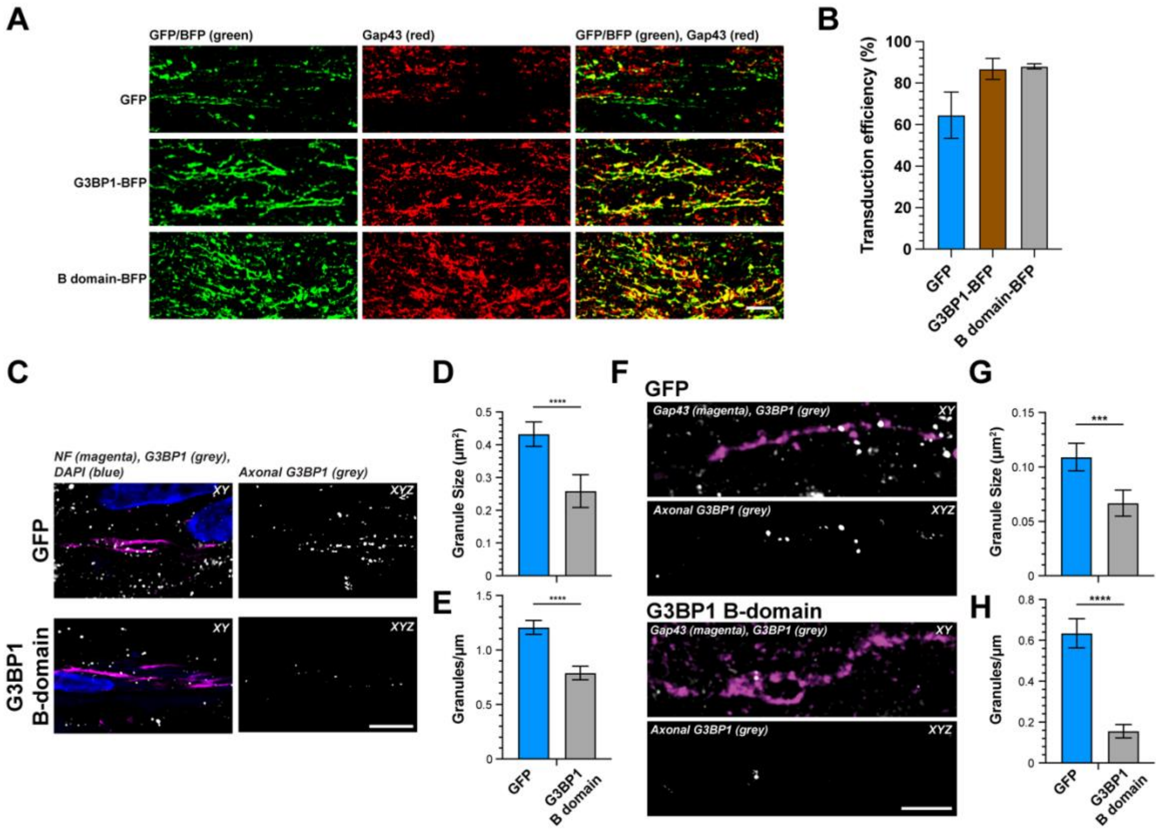


<u>Supplemental Figure S1</u>: ***G3BP1 B domain expression reduces G3BP1 granules in reticulospinal and optic nerve axons.***

***A-B,*** Representative exposure-matched GFP or BFP + GAP43 immunofluorescent images of crushed optic nerve are shown (**A**). Quantification of the percentage of GFP or BFP positive axons identified by GAP43 staining is shown as mean ± SEM (**B**, N ≥ 5 animals) [scale bar = 5 µm].

***C,*** Representative exposure-matched confocal images of reticulospinal axons regenerating in PNG as in Figure 1B with signals for neurofilament (NF) and G3PB1 immunoreactivity shown for animals expressing GFP vs. G3BP1 B-domain BFP. The left panel of each image pair shows merged signals in a single XY plane, and the right panel of each image pair shows XYZ projection of G3BP1 signals that overlap with NF across individual Z sections. Imaging sequences for G3BP1 were optimized to detect only the granular form of the protein [Scale bar = 10 μm].

***D,*** Diameter of axonal G3BP1 granules shown as mean ± SEM for reticulospinal axons as in A (N ≥ 116 granules over three repetitions for each condition; ***p ≤ 0.001 by Student's t-test).

***E,*** Density of axonal G3BP1 granules in optic nerve axons from sections as in A shown as mean ± SEM (N = 37 axons over three repetitions for each condition; ****p ≤ 0.0001 by Student's t-test).

***F,*** Representative exposure-matched confocal images of optic nerve axons as in Figure 1E with signals for GAP43 and G3PB1 immunoreactivity shown for animals expressing GFP vs. G3BP1 B-domain BFP. The left panel of each image pair shows merged signals in a single XY plane, and the right panel of each image pair shows XYZ projection of G3BP1 signals that overlap with GAP43 across individual Z sections [Scale bar = 10 μm].

***G,*** Mean size of axonal G3BP1 granules from optic nerves as in D shown as mean ± SEM (N ≥ 116 granules over three repetitions for each condition; ***p ≤ 0.001 by Student's t-test).

***H,*** Density of axonal G3BP1 granules in optic nerve axons from sections as in D shown as mean ± SEM (N = 37 axons over three repetitions for each condition; ****p ≤ 0.0001 by Student's t-test).

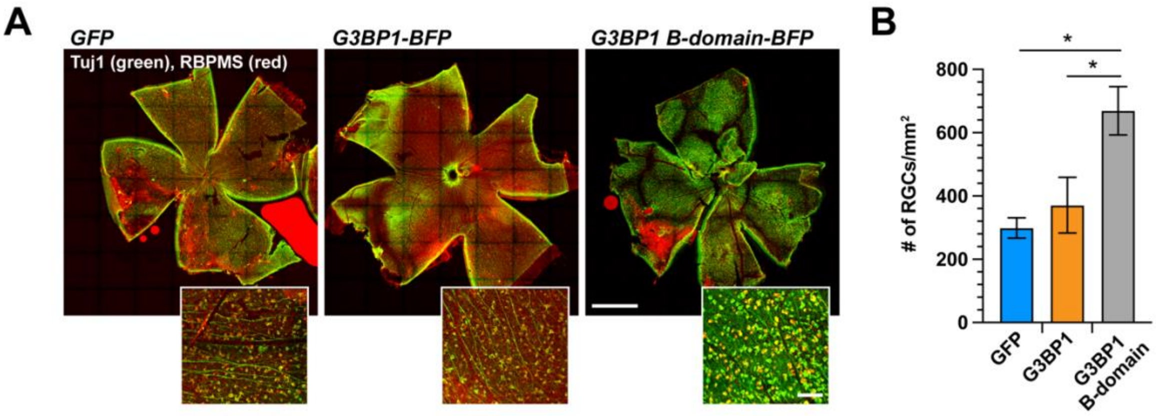


<u>Supplemental Figure S2</u>: ***Expression of G3BP1 acidic domain increases RGC survival after optic nerve injury.***

***A,*** Representative exposure-matched epifluorescence montage images for βIII Tubulin (Tuj1) and RBPMS stained retinae to identify RGCs from animals transduced with AAV2-GFP, -G3BP1-BFP, or G3BP1 B-domain-BFP shown. Inset panels show higher magnification [scale bars = 2 mm in large images and 100 µm in insets].

***B,*** Quantitation of surviving RGCs based on RBPMS positivity as in panel A shown as mean ± SEM ($N \geq 5$ animals; *$p \leq 0.05$ by one-way ANOVA with Tukey HSD post-hoc).

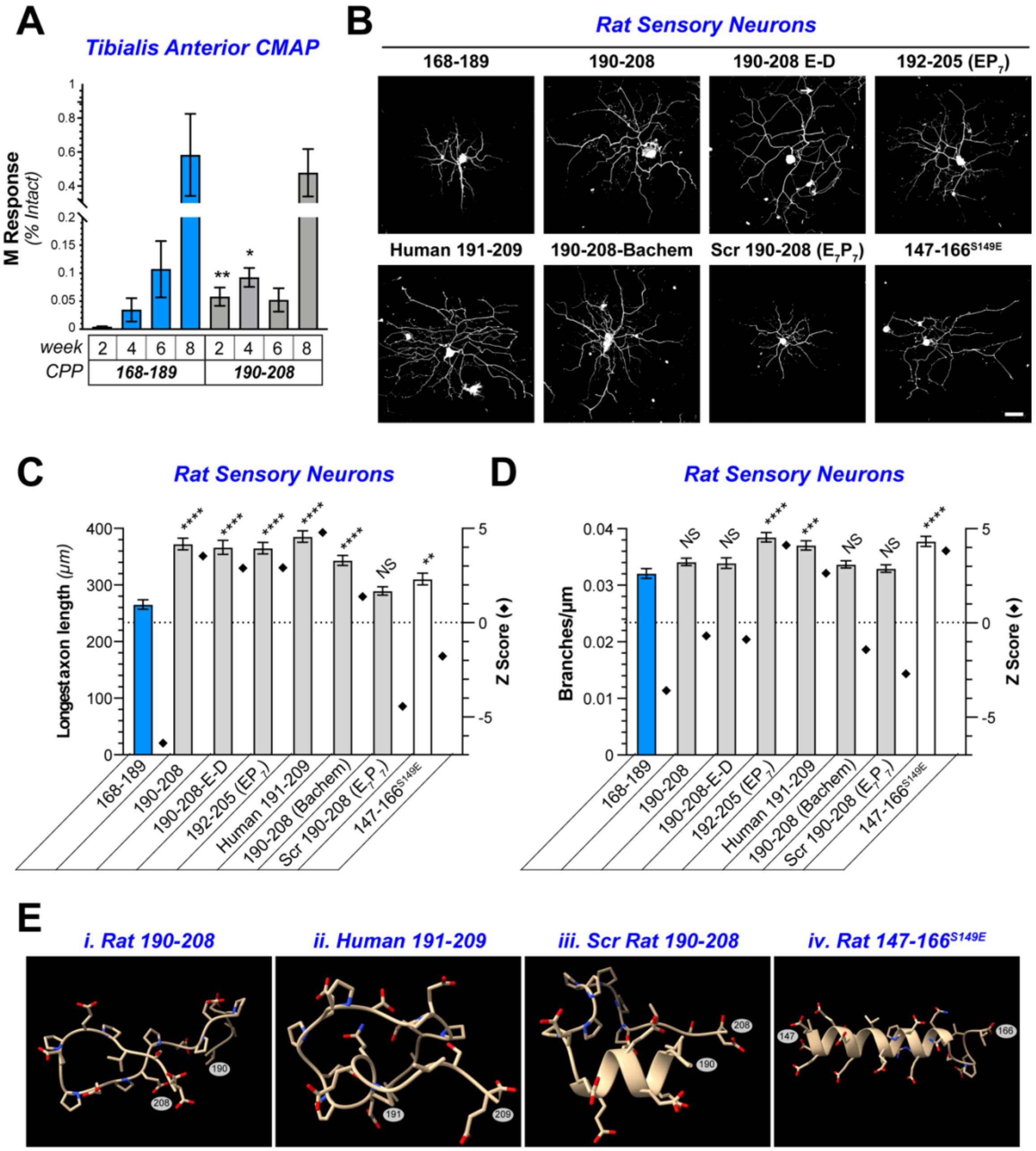


**Supplemental Figure S3**: ***G3BP1 190-208 cell permeable peptide promotes axon growth.***

***A,*** CMAPs for tibialis anterior muscle shown at indicated times post nerve crush injury for animals treated with G3BP1 168-189 vs. 190-208 CPP as average % intact M responses ± SEM (N ≥ 4 animals; *p ≤ 0.05 and **p ≤ 0.01 by Student's t-test for indicated data pairs).

***B,*** Representative exposure-matched epifluorescence images for NF immunostained DRG neurons treated with indicated peptides shown [scale bar = 100 µm].

***C-D,*** Quantitations of longest axon per neuron (**C**) and axon branching (**D**) for adult DRG neurons cultured on laminin as in B and treated with indicated G3BP1 CPPs and variants shown as mean ± SEM in (left Y-axes) and Z score vs. population mean (right Y-axes; N ≥ 352 neurons from three biological replicates; ** p ≤ 0.01, *** p ≤ 0.001, **** p ≤ 0.0001 by one-way ANOVA with Dunnett post-hoc).

***E,*** Predicted tertiary structures of rat G3BP1 190-208, human G3BP1 191-209, scrambled rat G3BP1 190-208 and rat G3BP1 147-166$^{S149E}$ peptide sequences from *PepFold3* are shown with N- and C-terminal residues indicated. Structures for the non-functional scrambled rat G3BP1 190-208 and rat G3BP1 147-166$^{S149E}$ peptide are quite distinct from the growth-promoting rat G3BP1 190-208, human G3BP1 191-209.

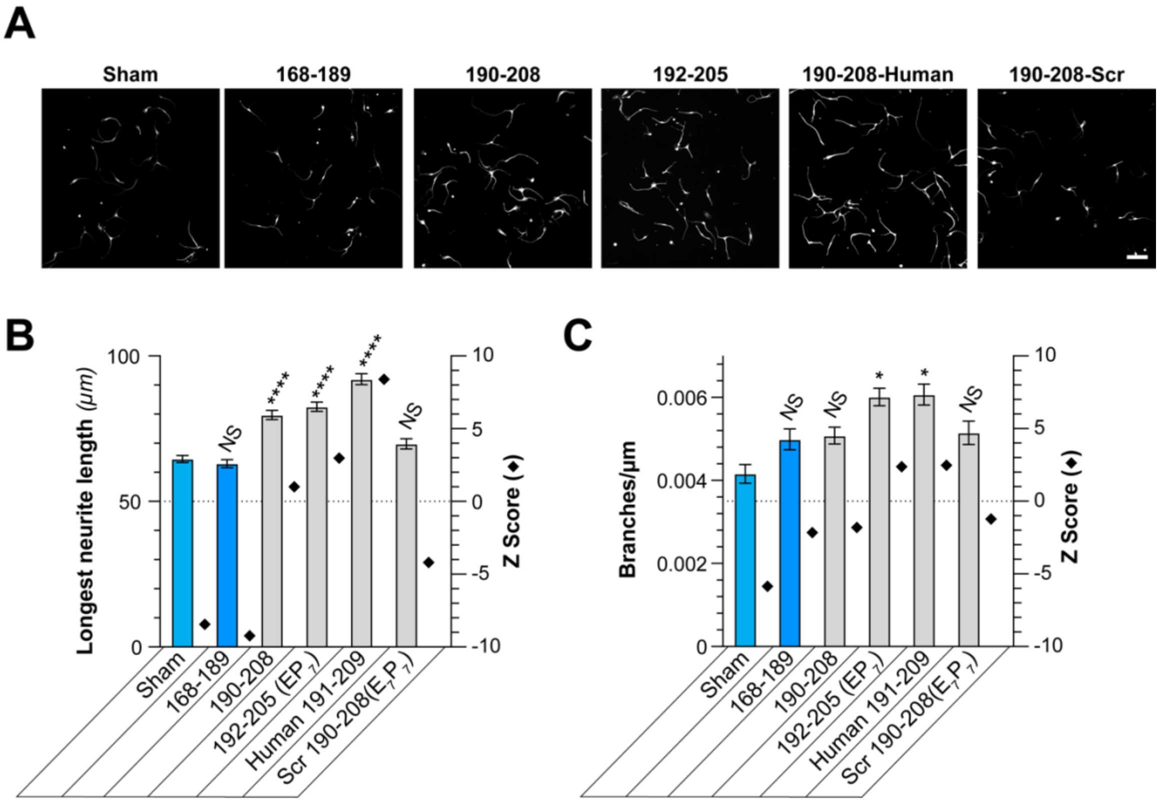


<u>Supplemental Figure S4</u>: ***Rat and human G3BP1 cell permeable peptides promote axon growth from human iPSC-derived glutamatergic neurons.***

***A,*** Exposure-matched epifluorescence images for βIII tubulin-immunostained hiPSC-derived glutamatergic neurons untreated (Sham) or treated with indicated CPP [scale bar = 100 µm].

***B-C,*** Quantitations of longest neurite per neuron (**B**) and neurite branching density (**C**) in hiPSC-derived glutamatergic neurons treated as in A shown as mean ± SEM (left Y-axes) and Z score vs. population mean (right Y-axes; N ≥ 1165 neurons across three biological replicates for each condition; * $p \leq 0.05$, **** $p \leq 0.0001$ vs. Sham by one-way ANOVA with Tukey HSD post-hoc).

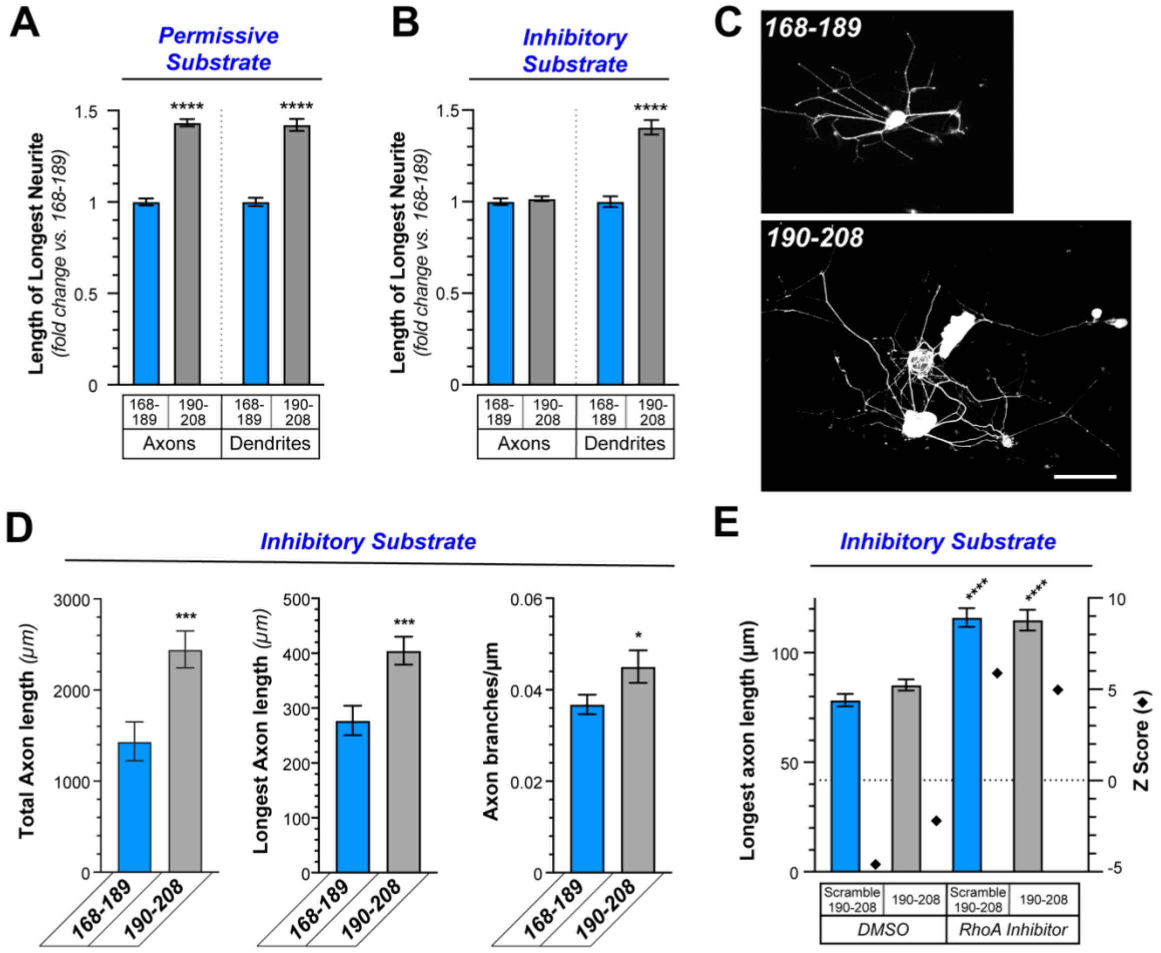


**Supplemental Figure S5**: ***G3BP1 cell permeable peptide increases axon branching on growth-inhibitory substrate.***

***A-B,*** Quantitations of longest distance axon or dendrite extends from each E18 cortical neurons in culture treated with G3BP1 168-189 or 190-208 CPP on laminin (**A**) or laminin + aggrecan (**B**) substrates shown as mean ± SEM (N ≥ 1042 neurons across 3 culture preparations for each condition; **** p ≤ 0.0001 by students T-test).

***C,*** Representative epifluorescence images of NF-immunostained adult DRG neurons cultured on laminin + aggrecan and treated with G3BP1 168-189. vs. 190-208 CPP are shown [scale bar = 50 µm].

***D,*** Quantitations of total axon length, longest axon/neuron, and axon branching for adult DRG neurons cultured as in C shown as mean ± SEM (N ≥ 71 neurons across three repetitions for each condition; ****p ≤ 0.0001 by one-way ANOVA with Tukey HSD post-hoc).

***E,*** Quantitation of longest axon per neuron for E18 cortical neurons cultured on PDL + aggrecan and treated with G3BP1 190-209 vs. scrambled 190-208 CPP as in Figure 4C-D is shown as mean ± SEM (left Y-axis) and Z score vs. population mean (right Y axis; N ≥ 259 neurons across three repetitions for each condition; ****p ≤ 0.0001 for indicated treatment group vs. scrambled 190-208-DMSO by one-way ANOVA with Tukey HSD post-hoc).

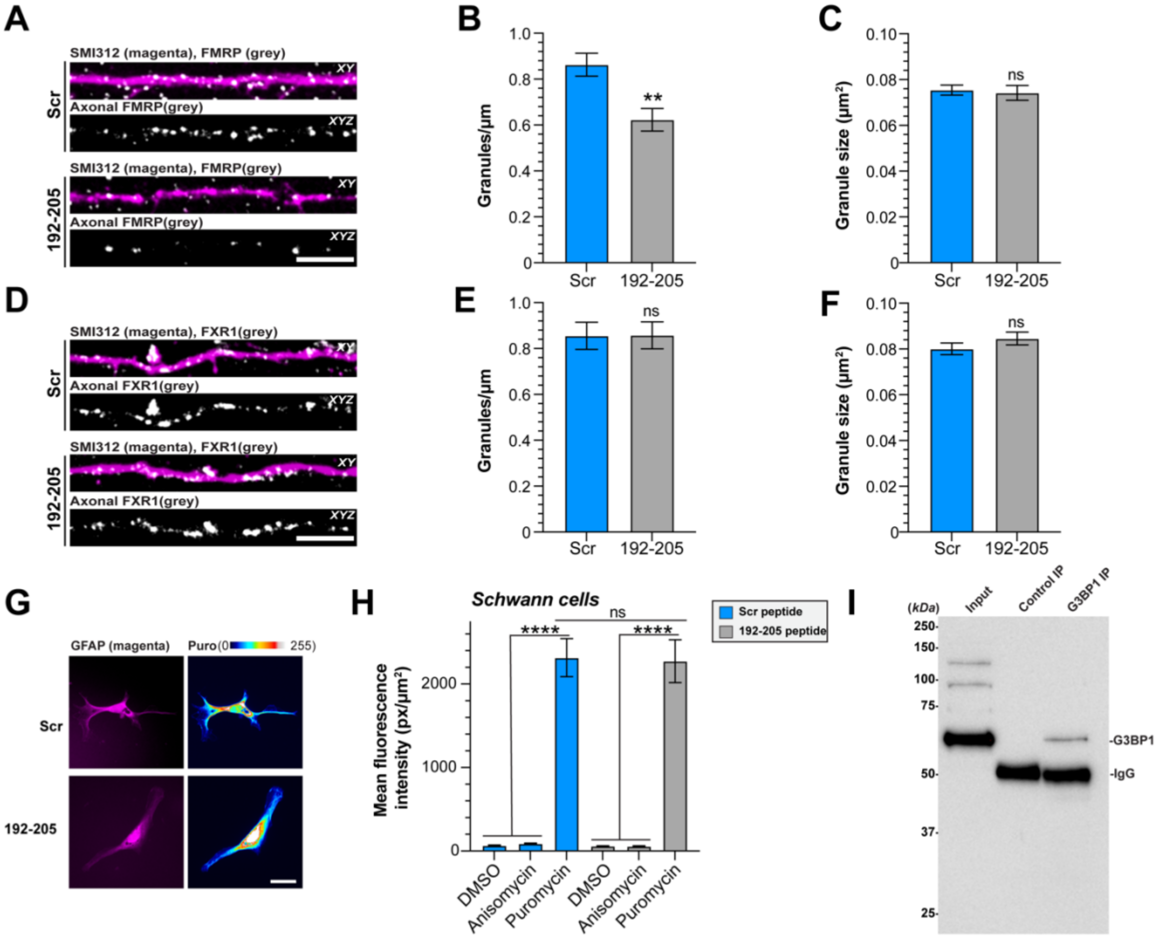


Supplemental Figure S6: ***G3BP1 granule disassembling CPP affects other stress granule proteins and does not activate glial protein synthesis.***

***A-C,*** Representative exposure-matched confocal images of axons of embryonic rat cortical neurons labeled with axonal neurofilament marker SMI312 and FMRP (**A**) are shown as in Figure 5A. Quantitation of the density (**B**) and size (**C**) of axonal FMRP granules shown as mean ± SEM (N = 36 axons across three biological replicates, $**p \leq 0.01$ by Unpaired Students t-test in B and $N \geq 484$ granules across three biological replicates, p values by Mann Whitney U test in C) [scale bar = 5 $\mu$m].

***D-F,*** Representative exposure-matched confocal images of axons of embryonic rat cortical neurons labeled with axonal neurofilament marker SMI312 and FXR1 (**D**) are shown as in A.

Quantitation of the density (**E**) and size (**F**) of axonal FXR1 granules shown as mean ± SEM (N = 36 axons across three biological replicates, p values by Unpaired Students t-test in E and N ≥ 911 granules across three biological replicates, p values by Mann Whitney U test in F) [scale bar = 5 μm].

***G-H,*** Exposure-matched epifluorescence images for GFAP + Puromycin immunostained adult rat Schwann cells after CPP treatment are shown (**G**). Quantitation of the amount of puromycin in adult rat Schwann cells after CPP treatment shown as mean ± SEM (**H**; N ≥ 35 Schwann cells across three biological replicates; p values by Kruskal Wallis test with Dunn's multiple comparisons) [scale bar = 25 μm].

***I,*** Western blot validating G3BP1 immunoprecipitation used in the G3BP1 RNA co-immunoprecipitation shown in Fig. 5J-M.
